# Inner membrane protein OutB is covalently attached to peptidoglycan in the γ-proteobacterium *Dickeya dadantii*

**DOI:** 10.1101/2024.09.03.610988

**Authors:** Xavier Nicolai, Yucheng Liang, Florence Ruaudel, Magdalena Narajczyk, Robert Czajkowski, Filippo Rusconi, Michel Arthur, Vladimir E. Shevchik

## Abstract

Gram-negative bacteria possess a multilayered envelope comprising an inner membrane (IM), a thin peptidoglycan (PG) layer, and an outer membrane (OM). In *Escherichia coli* and certain other γ-proteobacteria, Braun lipoprotein (Lpp) covalently tethers the OM to PG. Only a few other OM proteins have been found to be covalently linked to PG in Gram-negative bacteria. Here, we showed that in the phytopathogenic γ-proteobacterium *Dickeya dadantii*, an IM protein, OutB, is covalently attached to PG, thereby tethering itself and the associated type 2 secretion system to the cell wall. In contrast to Lpp, OutB reaches the PG layer from the IM side. By modifying the length of Lpp, which would displace the PG layer in the periplasm, we found that the elongated Lpp+21 improved OutB attachment to PG, whereas the shortened LppΔ21 reduced it. We showed that two L,D-transpeptidases, Ldt03 and Ldt84, tether Lpp and OutB to PG by the same catalytic mechanism involving the formation of an amide bond between their C-terminal lysine and the stem peptide. Ldt03 and Ldt84, each display substrate specificity for the type of peptide stem and preferentially cross-link Lpp to monomeric and dimeric muropeptide, respectively. The C-terminal Lpp-like box of OutB is almost identical to that of Lpp; it tolerates substantial amino acid substitutions and allows PG-tethering of a *bona fide* periplasmic protein. Thus, it seems possible that the repertoire of periplasmic and membrane proteins tethered to PG may be more extensive than currently assumed.

## Introduction

The envelope of Gram-negative bacteria consists of an inner membrane (IM) and an outer membrane (OM) that delineate the periplasm containing a thin layer of peptidoglycan (PG) (1). PG is an essential component of the cell envelope, which mechanically sustains the turgor pressure of the cytoplasm (2). The mesh-like structure of the PG macromolecule consists of glycan chains made of alternating *N*-acetylglucosamine (GlcNAc) and *N*-acetylmuramic acid (MurNAc) residues linked by β-1,4-bonds (3). The lactoyl group of MurNAc residues is linked via an amide bond to a pentapeptide stem, which in *Escherichia coli* consists of the sequence L-Ala^1^-D-Glu^2^-DAP^3^-D-Ala^4^-D-Ala^5^, wherein DAP is diaminopimelic acid (4). In this bacterium, the stem peptides are mainly cross-linked to each other via D-Ala^4^→DAP^3^ amide bonds (4→3 cross-links) formed by D,D-transpeptidases of the penicillin-binding protein (PBP) family (5, 6). In addition, L,D-transpeptidases form DAP^3^→DAP^3^ (3→3) cross-links, which account for 3% to 10% of the cross-links, depending on the growth phase (7, 8). In certain α- and β-proteobacteria, an alternative type of L,D-transpeptidase catalyzes the formation of 1→3-type cross-links between L-Ala^1^ and DAP^3^ (9, 10). Despite the different linking sites, 3→3 *versus* 1→3, both these Ldt families share a YkuD-like catalytic domain (PF03734). In addition, another class of 3→3 cross-linking Ldts possess a structurally unrelated VanW-type catalytic domain (PF04294) (11).

The attachment of the PG layer to both the IM and OM is essential to maintain the integrity of the cell envelope of Gram-negative bacteria. In the majority of Gracilicutes, two OM proteins, OmpA and Pal, non-covalently attach the OM to the PG via their conserved PG-binding domains (12–15). Pal is a component of the Tol-Pal trans-envelope complex that links the PG to both the IM and OM (16). Furthermore, in a subclade of γ-proteobacteria, including *E. coli*, the Braun lipoprotein (Lpp) covalently tethers the OM to the PG layer (12, 17). The N-terminal Cys residue of mature Lpp is acylated by three fatty acids, which are embedded in the inner leaflet of the OM. The side-chain ε-amino group of the C-terminal Lys residue of Lpp (Lys^58^) is covalently linked to the α-carbonyl of the DAP residue of PG tripeptide stems (18, 19). This cross-linking reaction is catalyzed by specialized L,D-transpeptidases, ErfK (LdtA), YbiS (LdtB), and YcfS (LdtC) (20). Three Lpp molecules form together a trimeric helix that tethers the OM to the PG layer (21). Lpp is the most abundant protein in *E. coli* (10^6^ copies per cell), about one-third of which is attached to PG, accounting for the covalent modification of about one in ten PG peptide stems (18, 22). For the past five decades, Lpp has remained the sole protein known to be covalently linked to PG in Gram-negative bacteria. However, two recent studies have shown that in some α- and γ-proteobacteria lacking any *lpp* ortholog, certain β-barrel outer membrane proteins (OMPs) are covalently linked to the DAP residue of the stem peptide via their N-terminal Ala or Gly residue (23, 24). These OMPs provide an alternative mechanism for maintaining envelope stability through the tethering of the OM to the PG. Moreover, in the γ-proteobacterium *Coxiella burnetii*, the OM lipoprotein LimB, a functional analog of Lpp, is attached to PG through an internal lysine residue (23). In addition to its osmoprotective role, the PG layer serves as a scaffold for the non-covalent attachment of various proteins and protein complexes. For instance, many trans-envelope machineries that span the cell envelope of Gram-negative bacteria are non-covalently linked to the cell wall by specialized PG-binding domains of various types, such as AMIN, LysM, SPOR, or OmpA-like (25–28).

*Dickeya dadantii* is a plant pathogenic γ-proteobacterium that secretes an array of virulence effectors via a type 2 secretion system (T2SS), termed Out (29). The Out system is a trans-envelope complex comprising fourteen proteins: OutB to OutM, OutO, and OutS (30). Among these proteins, OutB acts as a scaffolding protein facilitating the assembly of the OM pore formed by secretin OutD (31). OutB is anchored to the IM by its N-terminal transmembrane segment (TMS), followed by a 75-residue linker and the Homology Region (HR) (PF16537), ended by a 30-residue C-terminal extension (CTE) (Fig. 1A).

**Figure 1.**
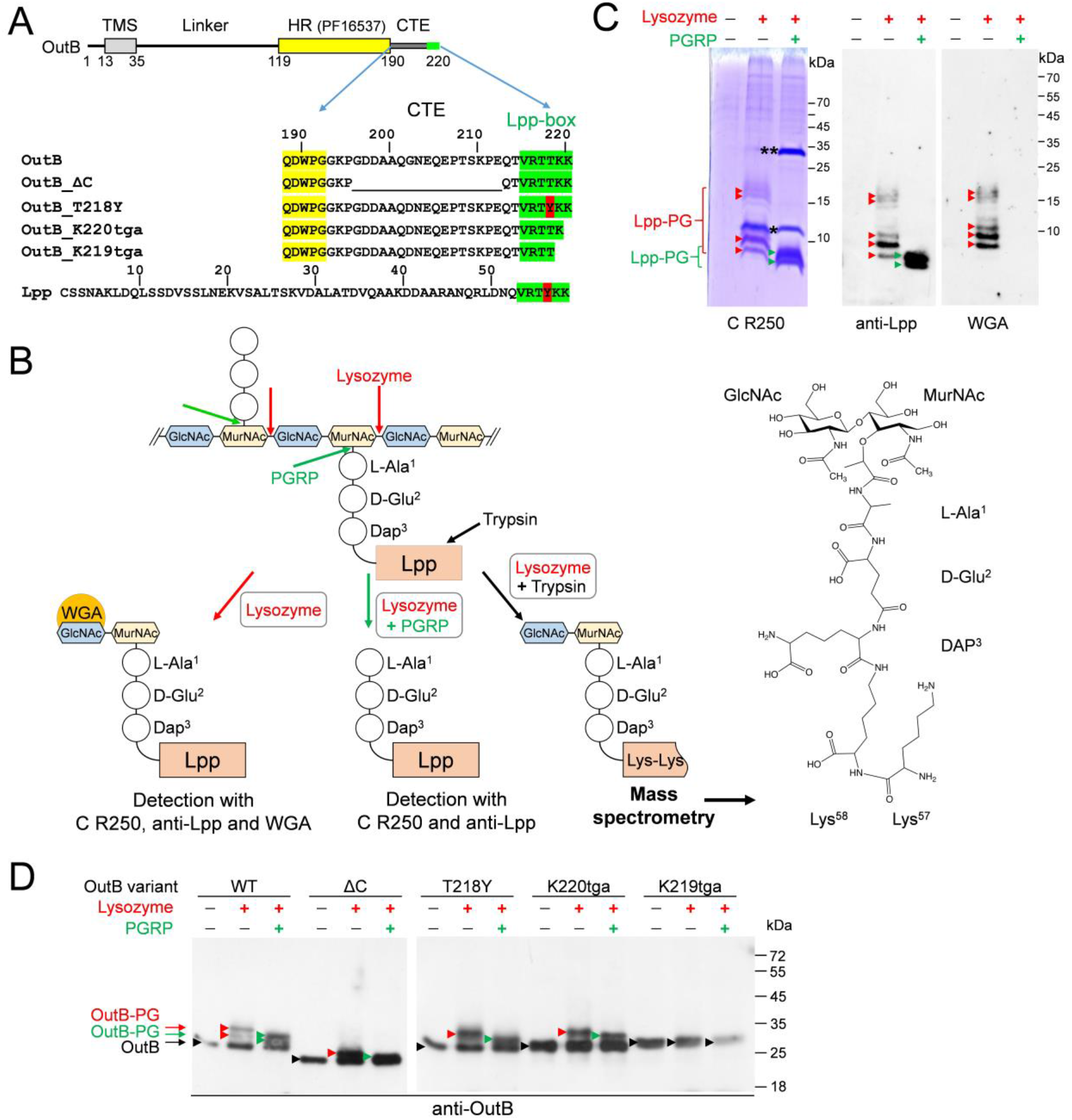
Covalent attachment of OutB and Lpp_Dda_ to PG in *D. dadantii*. (**A**) Domain organization of OutB. OutB comprises a transmembrane segment (TMS), a periplasmic linker, a homology region (HR) (in yellow), a C-terminal extension (CTE) carrying the Lpp box (in green). OutB variants carrying mutations in the CTE or Lpp-box are aligned together with the mature Lpp of *D. dadantii*. Sequence similarity between OutB and Lpp_Dd_ is limited to the Lpp box. (**B**) Schematic of the analysis of PG purified from *D. dadantii*. The cleavage sites of lysozyme, PGRP and trypsin are shown with red, green, and black arrows, respectively. Abbreviations: GlcNAc, *N*-acetylglucosamine; MurNAc, *N*-acetylmuramic acid; DAP, diaminopimelic acid. Wheat germ agglutinin (WGA) recognizes GlcNAc residues. Structure of muropeptide from *D. dadantii* PG attached to two C-terminal Lys residues as determined by mass spectrometry. (**C**) SDS-PAGE and Western blot analyses of the protein content of *D. dadantii* PG. PG was digested or not with lysozyme and PGRP amidase as indicated. Lpp-PG adducts attached to muropeptides were detected with Coomassie R250 (C R250) (left panel), anti-Lpp antibodies (middle panel), or WGA (right panel). Lpp-PG adducts generated by lysozyme and PGRP are shown with red and green arrowheads, respectively. The positions of lysozyme and PGRP are indicated by one and two asterisks, respectively. (**D**) Western blots of PG preparations purified from *D. dadantii* expressing the indicated OutB variants (shown in panel A). PG was digested or not with lysozyme and PGRP amidase and probed with anti-OutB antibodies. OutB-PG adducts generated by lysozyme and PGRP are indicated with red and green arrowheads, respectively. The position of “free” OutB forms is shown with black arrowheads.

Here, we employed a combination of biochemical, genetic, and mass spectrometry (MS) analyses to demonstrate that OutB possesses a genuine Lpp-like box and that its C-terminal Lys residue is covalently attached to the stem peptide of the *D. dadantii* PG by the same catalytic mechanism as that of Lpp. Two L,D-transpeptidases, Ldt03 and Ldt84, were identified as being involved in this process, with a preference for attaching the protein to muropeptide monomers and 4→3 cross-linked dimers, respectively. To determine how spatial constraints across the periplasm control the attachment of OutB to PG, we generated Lpp variants of various length, which would displace the PG layer in the periplasm. We find that the elongated Lpp+21 enhances OutB attachment to PG, whereas the shortened LppΔ21 has the opposite effect, reducing it. Mutagenesis analysis of the Lpp-box showed that it tolerates substantial amino acid substitutions and enables the attachment of a periplasmic reporter protein to PG. Taken together, these results suggest that certain other periplasmic and/or membrane proteins exposed to the periplasm with a C-terminal Lys residue may be covalently attached to the PG in *D. dadantii* and potentially in other Gram-negative bacteria.

## Results

### Lpp_Dd_ is covalently linked to the *D. dadantii* PG

Our previous study indicated that the C-terminal extension (CTE) of OutB might bind PG (31). The CTE has no obvious similarity to any known family of PG-binding domains, yet five of the six C-terminal residues of OutB (VRTTKK) are identical to those of *D. dadantii* Lpp, Lpp_Dd_ (VRTYKK) (Fig. 1A). This prompted us to examine whether OutB is covalently attached to the PG layer, as would be the case for Lpp_Dd_. To this end, the PG of *D. dadantii* was purified by the hot SDS procedure, which potentially eliminates all the proteins except those either covalently linked to the PG or trapped within the PG sacculi. The PG was then digested with lysozyme, a muramidase that cleaves the MurNAc-GlcNAc β-1,4 bonds (Fig. 1B), thereby releasing disaccharide-peptide PG fragments, including those covalently linked to Lpp or, potentially, to OutB. Accordingly, SDS-PAGE and immunoblotting of this PG sample revealed a series of protein bands reactive with anti-Lpp antibodies (Fig. 1C, left and middle panels). Most of these Lpp species were also detected with the wheat germ agglutinin (WGA) (Fig. 1C, right panel), which recognizes GlcNAc residues (32), thereby indicating the presence of Lpp covalently linked to muropeptides. Consistent with this assumption, digestion of the PG with PGRP, an amidase from the weevil *Sitophilus zeamais* that cleaves the MurNAc-L-Ala^1^ amide bond (Fig. 1B) (33), resulted in the complete loss of the WGA-reactive species (Fig. 1C, right panel). Instead, the digestion with PGRP produced two higher-mobility Lpp adducts, involving only stem peptide moieties (Fig. 1C). These results suggest that LppDd is covalently linked to two distinct types of stem peptides. This hypothesis will be addressed below.

### OutB is covalently linked to the *D. dadantii* PG

The PG preparations used above to identify Lpp-muropeptide adducts were subsequently probed by immunoblotting for the presence of OutB covalently linked to PG (Fig. 1D, left panel). Notably, a certain quantity of OutB was detected in the intact, undigested PG (Fig. 1D, lane without addition of lysozyme or PGRP). This was tentatively attributed to the “free” unbound form of OutB that remained entrapped in the intact sacculi during PG extraction, but escaped from them during SDS-PAGE. Such a “free” form was not observed with Lpp (Fig. 1C), which has a much smaller size (58 *versus* 220 residues) and, in contrast to OutB, is naturally located outside the sacculi. Digestion of the PG with lysozyme produced two additional slower-migrating OutB species that could correspond to OutB linked to muropeptides (Fig. 1D, left panel). Accordingly, as observed above for Lpp, additional digestion with PGRP, which removes saccharide moieties from stem peptides, resulted in OutB adducts with slightly increased mobility (Fig. 1D, left panel). These species could correspond to OutB linked to peptide moieties of PG.

High background noise generated by very abundant Lpp-linked muropeptides impeded the detection of muropeptide-linked OutB adducts by WGA. To address this issue, PG was extracted from *D. dadantii Δlpp* mutant ectopically expressing *outB*. WGA-reactive OutB species was detected in this PG digested with lysozyme (Fig. S1), indicating the presence of GlcNAc-containing muropeptides linked to OutB. Collectively, these results show that OutB is covalently linked to the peptide moieties of PG.

### Lpp is attached to PG via a DAP→Lys cross-link

To characterize the molecular link connecting Lpp_Dd_ and OutB to PG, the structure of the *D. dadantii* PG was determined by mass spectrometry (MS). The chemical structure of the *D. dadantii* PG was found to be identical to that of *E. coli* (Fig. S2). Briefly, the main monomers consisted of the disaccharide GlcNAc-MurNAc and a tripeptide (L-Ala^1^-D-Glu^2^-DAP^3^) or a tetrapeptide (L-Ala^1^-D-Glu^2^-DAP^3^-D-Ala^4^) stem. The main dimers contained a tetrapeptide stem linked to a tripeptide (Tetra→Tri dimer) or to a tetrapeptide (Tetra→Tetra dimer) stem by a D-Ala^4^→Dap^3^ 4→3 cross-link formed by the D,D-transpeptidase activity of PBPs. A disaccharide-tripeptide monomer substituted by a Lys-Lys dipeptide was detected. This muropeptide corresponds to the product of partial protein digestion by trypsin leaving the two C-terminal Lys residues bound to a tripeptide stem (Fig. 1B). A 4→3 cross-linked dimer substituted by two Lys residues (Tetra→Tri→Lys-Lys) was also detected (Fig. S2). This muropeptide may originate from the attachment of a protein to a 4→3 cross-linked dimer or from the cross-linking of a disaccharide-peptide to a preformed Tri→protein adduct. Additionally, minor quantities of muropeptides comprising a single Lys residue were detected, indicative of partial cleavage of the Lys-Lys peptide bond by trypsin. Since Lpp_Dd_ and OutB both share the identical Lys-Lys C-terminal sequence (Fig. 1A), the Tri→Lys-Lys and Tetra→Tri→Lys-Lys molecular species detected by MS may potentially originate from the digestion of both PG-bound Lpp_Dd_ and OutB adducts by trypsin.

To investigate this possibility, we conducted a PG analysis of *D. dadantii Δlpp* mutant. The Tri→Lys-Lys and Tetra→Tri→Lys-Lys were not detected in the PG extracted from this mutant. This result indicates that, in contrast to immunoblotting (Fig. S1), the sensitivity of the MS analysis is sufficient only to detect muropeptide-linked adducts derived from the highly abundant Lpp_Dd_ protein.

### The Lpp box of OutB is necessary and sufficient for the covalent attachment to PG

The 6-residue Lpp-like box of OutB is located at the C-terminal end of the CTE and is preceded by a 22-residue region following the HR domain (Fig. 1A). This part of CTE is rich in charged residues and shares some features with sugar-binding motifs (34). However, removal of a substantial portion of the CTE outside the Lpp-box (residues G^196^ to E^212^) did not prevent covalent attachment of the resulting OutBΔC to PG (Fig. 1A and 1D), indicating that the deleted region is not essential for this purpose.

To further address the issue, either the full-length 30-residue C-terminal extension of OutB or only its truncated 13-residue portion (as in OutBΔC) was fused to the C-terminus of β-lactamase BlaM, a *bona fide* periplasmic protein, thereby forming Bla-CTE and Bla-CTEΔC, respectively (Fig. 2A). Immunoblotting showed that both hybrids are covalently linked to PG (Fig. 2B). Indeed, digestion of the corresponding PG with lysozyme yielded three BlaM-reactive adducts of lower electrophoretic mobility, consistent with the presence of covalently attached muropeptides. Addition of PGRP amidase generated two Bla species of a smaller apparent size, indicating elimination of glycan moieties (Fig. 2B). These data show that the Lpp-box of OutB constitute a genuine PG-linking motif that allows covalent attachment of the periplasmic reporter protein to the *D. dadantii* PG.

**Figure 2.**
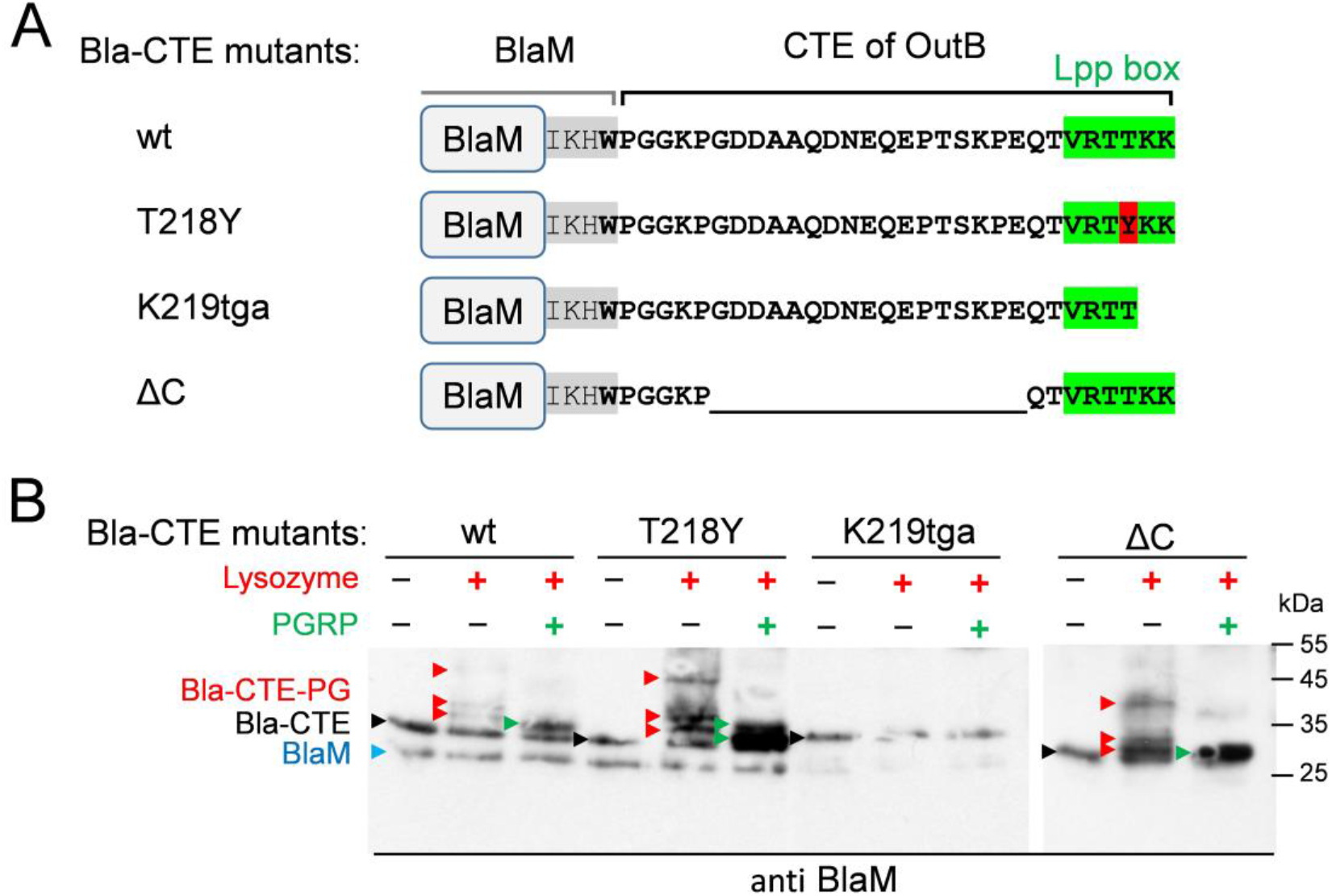
Lpp-box of OutB allows attachment of the β-lactamase BlaM to PG. (**A**) Schematic of the BlaM-CTE fusions. The C-terminus of the native BlaM is highlighted in grey; the rest of the sequence corresponds to the CTE of OutB (residues P191 to K220). The Lpp-like box is highlighted in green and the T218Y substitution is in red. The numbering of residues substituted in BlaM-CTE mutants is that of the OutB CTE. (**B**) Western blot analysis of PG purified from *D. dadantii* expressing the indicated Bla-CTE variants. PG was digested or not with lysozyme and PGRP amidase and probed with anti-BlaM antibodies. The positions of BlaM and “free” forms of Bla-CTE fusions are shown with blue and black arrowheads, respectively. Bla-CTE-PG adducts generated by lysozyme and PGRP are indicated with red and green arrowheads, respectively.

### Mutagenesis of the Lpp box

Unlike *E. coli* Lpp, which carries a C-terminal Arg-Lys pair, the C-termini of Lpp_Dd_ and OutB contain two Lys residues (Fig. 1A). To evaluate the role of these residues in the attachment of OutB to PG, an Opal stop codon was introduced in place of the codons encoding Lys^219^ or Lys^220^. As expected, deletion of the two C-terminal Lys abolished the attachment of the resulting OutB_K219tga to PG (Fig. 1D). However, OutB_K220tga, which lacks the C-terminal Lys^220^, was tethered to PG at the wild-type level (Fig. 1D). This suggests that the remaining penultimate Lys^219^, which has become the C-terminal residue, may link OutB to PG. Strikingly, the equivalent substitution in Lpp (loss of the C-terminal Lys^58^) showed about 50-fold reduction in the amount of PG-linked LppΔ58K (Fig. S3A and S3B). Accordingly, LppΔ58K was barely active in the SDS susceptibility assay with the *D. dadantii Δlpp* mutant (Fig. S3C). These results show that a single Lys residue at the C-terminus of the Lpp box is both necessary and sufficient for covalent attachment to PG, but its reactivity varies depending on the protein.

The sole difference between the Lpp-box of OutB and Lpp_Dd_ is the presence of Thr *versus* Tyr at the antepenultimate position (Fig. 1A). The above data indicated that the PG-linking activity of LppΔ58K is more drastically affected than that of OutB_K220tga. To render the truncated Lpp-box of LppΔ58K identical to that of OutB_K220tga (VRTYK *versus* VRTTK), an Y56T substitution was introduced into LppΔ58K. This did not improve the attachment of the resulting LppY56T/Δ58K to PG or its activity in the SDS susceptibility assay with the *D. dadantii Δlpp* mutant (Fig. S3). More markedly, the Y56T substitution in the full-length Lpp notably reduced the attachment of the resulting LppY56T to PG and diminished its efficacy in the SDS susceptibility assay (Fig. S3). In contrast, the reciprocal T218Y substitution in OutB or in the Bla-CTE hybrid appeared to enhance the attachment of the proteins to PG (Fig. 1D and 2B). Thus, the polymorphism of the antepenultimate position, Thr (OutB) *versus* Tyr (Lpp_Dd_), has some effect on the tethering of these proteins to PG. Taken together, these results show that the Lpp-box can tolerate substantial amino acid substitutions, depending on the tethered protein.

### Anchoring of OutB to the inner membrane is not essential for its linkage to PG

The N-terminal IM anchor is critical for the scaffolding function of OutB towards the secretin OutD (31). To assess the role of this transmembrane segment (TMS) in the covalent attachment of OutB to PG, a soluble periplasmic variant, OutB_SP, was generated, in which the TMS was replaced by a cleavable signal peptide (Fig. S4A). In contrast to the full-length OutB and other OutB variants, no “free” form of OutB_SP was detected in the intact, undigested PG (Fig. S4B), indicating that OutB_SP is too small (∼19 kDa) to be retained by the PG meshwork. Lysozyme digestion of this PG sample yielded an anti-OutB reactive adduct of ∼30 kDa, whereas the addition of PGRP amidase resulted in a notable reduction in its amount and the appearance of a 25 kDa species (Fig. S4B). These data show that the anchoring of OutB to the IM is not mandatory for the tethering to the PG.

### Alteration of the Lpp_Dd_ length affects the integrity of the *D. dadantii* envelope

In *E. coli* and some other γ-proteobacteria, Lpp tethers the PG layer to the OM, thereby controlling the size of the periplasm. Altering the length of Lpp has been shown to alter the width of the periplasm and the distance between the PG layer and each of the two cell membranes (35–37). We wondered whether such variations in the Lpp_Dd_ length would affect the attachment of OutB to PG. At the same time, we tested whether alterations in the length of OutB could also affect its linkage to PG (Fig. 3A).

**Figure 3.**
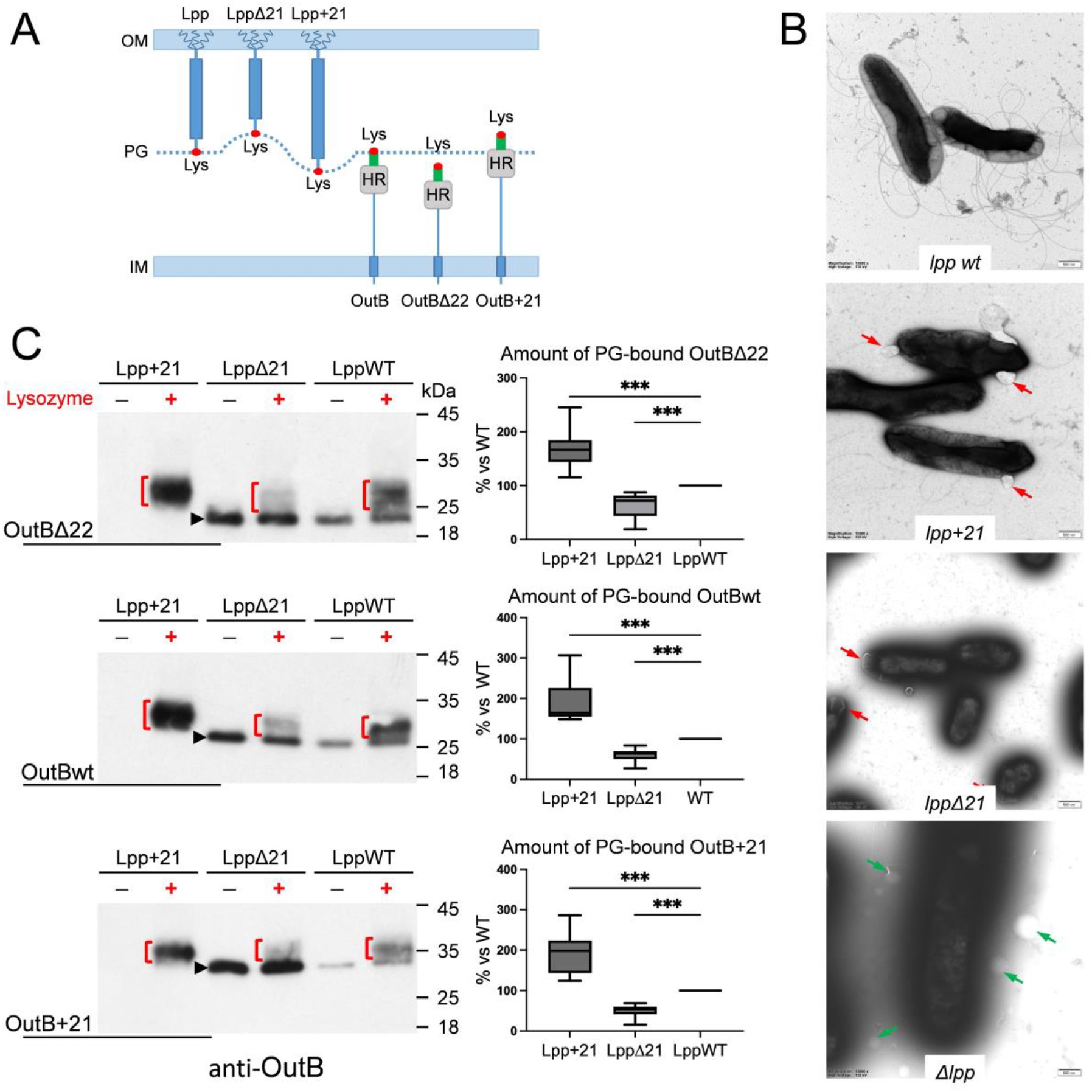
Altering the Lpp_Dd_ length affects OutB attachment to PG. (**A**) Schematic of the Lpp and OutB length variants used in the study (see Fig. S5A and S8A for details). (**B**) Negative stain TEM of the indicated *D. dadantii lpp* mutant strains. Blebs formed by *lppΔ21* and *lpp+21* cells are shown with red arrows and vesicles around *Δlpp* mutant, with green arrows. (**C**) OutBΔ22, OutBwt and OutB+21 length variants (indicated on the left) were expressed in *D. dadantii lpp* length mutants (*lppWT, lppΔ21* and *lpp+21*, indicated on the top of Western blots). PG was extracted from these nine strains, digested or not with lysozyme and probed with anti-OutB. OutB-PG adducts generated by lysozyme are shown with red brackets. “Free” forms of OutB variants are indicated with black arrowheads. The Westerns blots are representative of five technical replicates of two biological experiments. Box-plots show relative amounts of muropeptide-linked adducts for each OutB length variant in each *D. dadantii lpp* length mutant (shown with red brackets in the corresponding Western blots). The data were analyzed with PRISM software using two-sample Mann-Whitney test by comparing the values in *lpp+21* and *lppΔ21* mutants to the wild-type strain considered as 100%. Median, quartiles and whiskers are shown. *** denote statistically significant differences with *P* values <0,001.

In this aim, we constructed *D. dadantii* mutants producing a shortened or a lengthened Lpp_Dd_ variant, LppΔ21 and Lpp+21, respectively. The *lppΔ21* and *lpp+21* alleles were introduced into the *D. dadantii* chromosome in place of the wild-type *lpp* gene (Fig. S5A and S5B). Analysis of PG from these strains revealed that the apparent size of the LppΔ21 and Lpp+21 species linked to muropeptides varied according to the length of each variant (Fig. S5C). The *lppΔ21* and *lpp+21* mutants exhibited a range of phenotypic defects, which were, however, strain-specific. For instance, *D. dadantii lppΔ21* formed very mucoid colonies and was non-motile in swimming and swarming assays, but had wild-type resistance to SDS (Fig. S5D and S5E). In contrast, *D. dadantii lpp+21* showed increased susceptibility to SDS, yet its motility was barely affected, and its colonies had a regular, wild-type appearance (Fig. S5D and S5E). Electron microscopy (EM) revealed striking alterations in the cell morphology of the mutant strains (Fig. 3B). *D. dadantii lppΔ21* cells were significantly shorter compared to the wild-type strain and the *lpp+21* mutant, and were surrounded by a black oriole in negatively stained EM (Fig. 3B and S6). Both the *lppΔ21* and *lpp+21* mutants produced a sort of blebs, which remained attached to the cells, unlike the vesicles in the *Δlpp* strain (Fig. 3B). In scanning EM, such envelope defects were visible as cavities at the cell poles of the *lppΔ21* mutant (Fig. S7). Some of these phenotypes have been previously observed in *E. coli* and *Salmonella enterica* mutants expressing Lpp-length variants (35–38).

### Increasing Lpp length improves OutB attachment to PG

Subsequently, to evaluate the impact of OutB and Lpp length alterations on OutB tethering to PG, we constructed OutB variants with altered length, OutBΔ22 and OutB+21 (Fig. 3A and S8A) and combined them with the Lpp-length variants. In this way, the *outB, outBΔ22*, and *outB+21* genes were ectopically expressed in the *D. dadantii* wild-type, *lppΔ21*, and *lpp+21* strains. The PG from these strains (nine combinations) was digested with lysozyme and analyzed by immunoblotting with anti-OutB (Fig. 3C and S8B). This analysis revealed that the quantity of OutB-PG species steadily increased as a function of Lpp length, from LppΔ21 to Lpp and to Lpp+21, irrespective of the length of OutB itself (OutBΔ22, OutBwt, or OutB+21) (Fig. 3C, S8B and 8C). Specifically, the highest amount of muropeptide-bound OutB variants was observed in *D. dadantii lpp+21*, while their lowest amount was detected in *D. dadantii lppΔ21*. These results suggest that the length of Lpp, which controls the width of the periplasm, is a critical factor influencing the accessibility of the Lpp-box of OutB to PG.

It is noteworthy that the “free” forms of OutB-length variants showed an inverse trend. Their levels gradually decreased as a function of Lpp length in the series of strains, from *lppΔ21* to *lpp* and to *lpp+21* (Fig. 3C). This observation indicates that in the presence of LppΔ21 and Lpp+21, the PG meshwork is, respectively, more and less densely fitted than with the wild-type Lpp.

### Lpp_Dd_ and OutB are covalently tethered to PG by two L,D-transpeptidases with partially redundant functions

Analysis of the *D. dadantii* 3937 genome (asap.genetics.wisc.edu) revealed the presence of four genes encoding putative L,D-transpeptidases, which carry a characteristic YkuD domain, ABF-0016403, ABF-0020084, ABF-0020070 and ABF-0046523, hereafter referred to as *ldt03, ldt84, ldt70* and *ldt23*, respectively (Fig. S9). To identify the Ldts that are able to link Lpp_Dd_ and OutB to the PG, we constructed *D. dadantii* mutants with single and double deletions of these genes. SDS-PAGE and western blot analyses of PG from these strains showed that the deletion of both *ldt03* and *ldt84* is necessary to suppress the attachment of Lpp_Dd_ and OutB to PG (Fig. 4). Accordingly, the loss of Lpp_Dd_ tethering to PG upon the deletion of both *ldt03* and *ldt84* resulted in an increased SDS susceptibility of the double mutant (Fig. S10B). Markedly, the residual SDS resistance of *D. dadantii Δldt03Δldt84* was notably higher than that of the *Δlpp* mutant, suggesting that the fraction of Lpp that is not covalently linked to PG plays a role in maintaining the integrity of the cell envelope.

**Figure 4.**
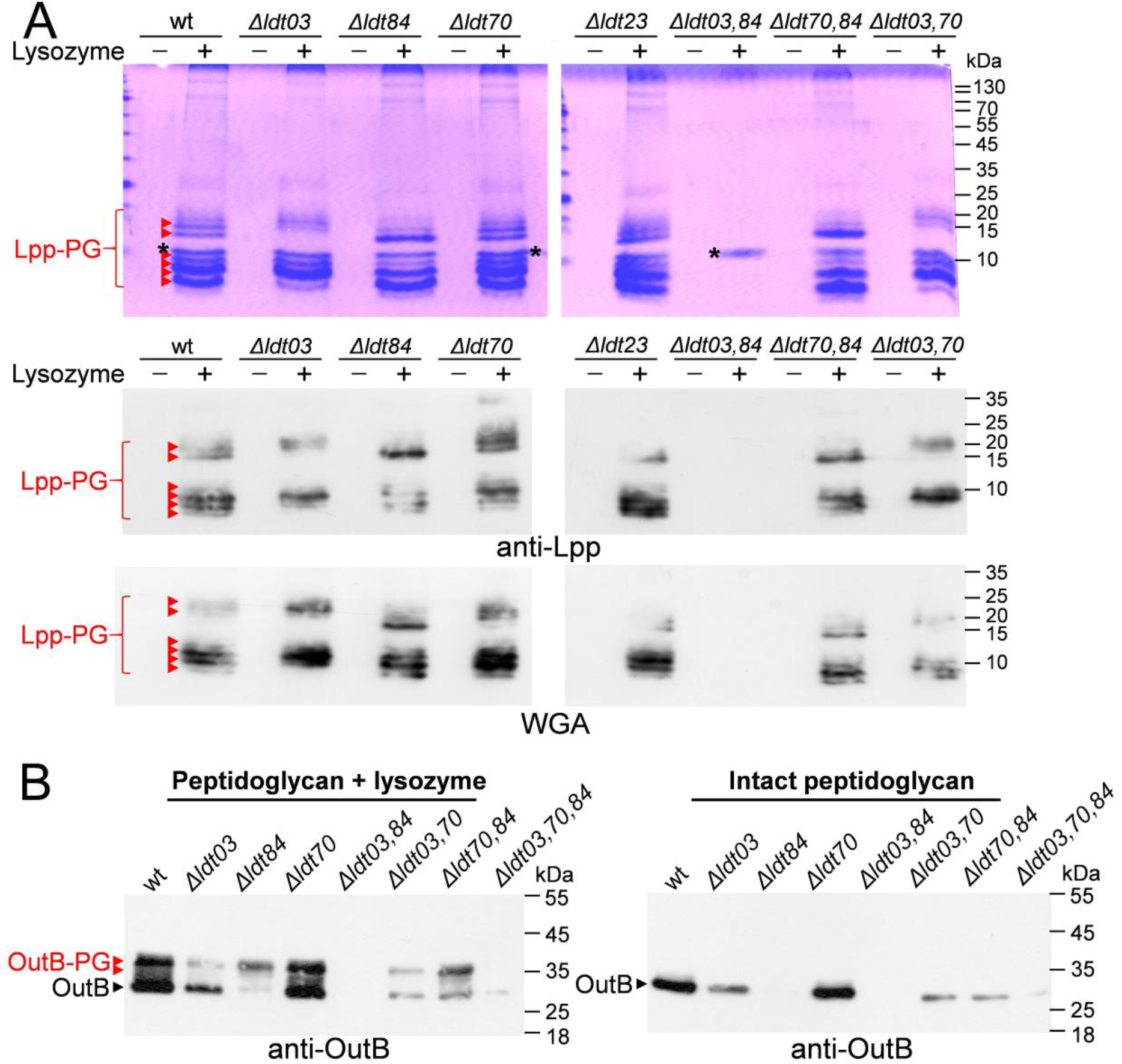
L,D-transpeptidases Ldt03 and Ldt84 are required for covalent attachment of Lpp and OutB to PG in *D. dadantii*. (**A**) SDS-PAGE and Western blot analyses of PG purified from the wild type (wt) *D. dadantii* and the indicated *Δldt* mutants. PG was digested or not with lysozyme and Lpp-PG species were detected with Coomassie R250 (upper panel), anti-Lpp antibodies (middle panel), or with WGA (lower panel); they are shown with red arrowheads. Position of lysozyme is indicated with an asterisk. Some other double and triple *Δldt* mutants are shown on Fig. S10. (**B**) The same PG samples as in panel A were probed with anti-OutB antibodies. “Free” form of OutB is shown with black arrowheads and OutB-PG adducts generated by lysozyme are indicated with red arrowheads.

Deletion of only *ldt03* or *ldt84* differently affected the abundance of various Lpp-PG and OutB-PG species (Fig. 4 and S10A), indicating that these enzymes exhibit certain degree of specificity for the PG-linking site. Consistent with this hypothesis, ectopic expression of *ldt03* in the double *Δldt03Δldt84* mutant resulted in an Lpp-PG pattern similar to that of the *Δldt84* mutant, and conversely, expression of *ldt84* yielded an Lpp-PG pattern similar to that of the *Δldt03* strain (Fig. S11A). Likewise, ectopic expression of either *ldt03* or *ldt84* in the wild-type *D. dadantii* was found to increase the amount of PG-bound OutB, yet more obviously with *ldt03* (Fig. S11B).

Deletion of the *ldt70* or *ldt23* genes, either alone or in combination with *ldt03* or *ldt84*, did not appear to affect the Lpp-PG and OutB-PG patterns or the SDS susceptibility of the corresponding mutants (Fig. 4A and S10). The catalytic site of Ldt23 is similar to that of LdtF (YafK), which catalyzes the cleavage of Lpp from the stem peptide in *E. coli* (Fig. S9) (39, 40). Ldt70 shows the same domain organization as LdtD (YcbB), which is responsible for the 3→3 cross-links of stem peptides (Fig. S9) (41). It seems plausible that Ldt23 and Ldt70 perform analogous functions to their respective *E. coli* orthologs.

### Ldt03 and Ldt84 display specificity for the nature of stem peptide

Given the notable dissimilarity between the Lpp-PG patterns produced by Ldt03 and Ldt84, we sought to elucidate the underlying molecular basis of these Ldt-specific patterns. In this order, MS analysis of PG from *D. dadantii Δldt03* and *Δldt84* mutant strains was performed. Muropeptides derived from PG were subjected to *rp*HPLC separation, and molecular species bearing the Lys-Lys moiety of Lpp were identified by MS (Fig. S12A). The relative abundance of each species was estimated based on their respective ion current intensities (Fig. S12B). This analysis revealed that the monomeric (Tri→Lys-Lys) muropeptide is predominant in the *D. dadantii Δldt84* mutant (Ldt03+), (79.1% ± 6.3%), whereas the dimeric (Tetra→Tri→Lys-Lys) muropeptide is predominant in the *D. dadantii Δldt03* mutant (Ldt84+) (72.0% ± 9.8%) (Fig. S12B). In agreement with this result, SDS-PAGE analysis of PGRP-digested PG from the *D. dadantii Δlpp Δldt03 Δldt84* triple mutant, which ectopically expresses *lpp-strep* with either *ldt03* or *ldt84*, resolved two Lpp-PG species consistent with those attached to monomeric and dimeric PG stem peptides (Fig. S12C). Once again, the putative monomer- and dimer-linked Lpp species were more abundant with Ldt03 and Ldt84, respectively. Taken together, these results show that Ldt03 and Ldt84 have a preference for tethering Lpp to monomeric and 4→3 cross-linked peptide stems, respectively.

### OutB is expressed in response to envelope stress associated with the plant infection process

OutB functions as a scaffolding protein that facilitates the assembly of the secretin OutD into the OM (31). The expression of *outD* is co-regulated with that of other genes of the *outC-outM* operon encoding the T2SS machinery, whereas the regulation of *outB* remains unclear (42, 43). To gain further insight, we examined the production of OutB in a series of *D. dadantii* regulatory mutants. Strikingly, high levels of OutB were observed in the *ΔphoP* and *ΔompR* mutants, but not in the *ΔkdgR* or *ΔpecS* mutants, as was the case with OutD (Fig. S13). The EnvZ/OmpR and PhoP/PhoQ systems are sensors of cell envelope and osmotic stresses (44). Consistent with this, OutB was massively produced at high osmolarity (0.2 M NaCl) (Fig. S13A). These results suggest that OutB may play a particularly important role in response to envelope stress. *D. dadantii* T2SS secretes a number of pectinases causing plant tissue maceration. Consistent with this, high levels of both OutB and OutD were detected in *D. dadantii* cells collected from rotted chicory leaves (Fig. S13B). These data show that during plant infection, OutB functions in concert with the cognate secretin OutD, reinforcing its attachment to the bacterial cell wall and thereby facilitating secretion of virulence factors by the T2SS.

## Discussion

In this study, we showed that the inner membrane protein OutB, a component of the *D. dadantii* T2SS, is covalently attached to PG by the same mechanism as Lpp. The site of attachment of OutB to the stem peptide of PG is located at the C-terminus of OutB and is nearly identical to that of Lpp. The originality of this finding lies in the substantial differences between OutB and a few other proteins known to be covalently linked to PG in Gram-negative bacteria. Indeed, Lpp and β-barrel OMPs are highly abundant proteins that tether the OM to the PG layer (19, 22–24). In contrast, OutB is an IM protein that binds the PG layer from an opposite side of the periplasm. Moreover, OutB is moderately expressed and does not appear to play a structural role in the cell envelope. Instead, OutB uses PG as a scaffold to better attach itself and the T2SS secretin pore to the cell wall (31).

We showed that two L,D-transpeptidases of *D. dadantii*, Ldt03 and Ldt84, catalyze the attachment of OutB and Lpp to PG. The majority of reported Ldts possess a YkuD catalytic domain (PF03734). The Ldts of the WanW family (PF04294) are much less prevalent and largely restricted to Gram-positive bacteria (11). No Ldt of the WanW family was identified in *D. dadantii*. Proteobacteria possess a variable number of YkuD-type L,D-transpeptidases, ranging from only one in *Neisseria gonorrhoeae and Helicobacter pylori* to 10 and 14 in *Legionella pneumophila* and *Agrobacterium tumefaciens*, respectively (23, 45). The *E. coli* genome encodes six proteins with a YkuD-type catalytic domain. Three of these enzymes, ErfK (LdtA), YbiS (LdtB) and YcfS (LdtC), are responsible for the tethering of Lpp to PG (20). The presence of multiple Ldts involved in the attachment of Lpp to PG may be attributed to their distinctive enzymatic properties and/or regulatory mechanisms. We provide evidence that the two *D. dadantii* L,D-transpeptidases, Ldt03 and Ldt84, which attach Lpp and OutB to PG, display a clear preference for specific types of muropeptides, monomeric and dimeric peptide stems, respectively. To the best of our knowledge, this study is the first demonstration of Ldt specificity for a particular type of peptide stem.

In comparison with a few other known proteins covalently linked to the PG of Gram-negative bacteria, OutB adopts a rather unusual topology. In OutB, the Lpp-box is grafted onto the periplasmic HR domain, which in turn is connected to the IM anchor by a 75-residue linker. The length of this linker varies even among closely related *Dickeya* species. For example, it is 83 residues in *D. dadantii subsp. dieffenbachiae* and 61 residues in *D. solani*. Consistent with this, we showed that a moderate truncation or extension of this linker (22 and 21 residues, respectively) has no discernible impact on OutB attachment to PG. In contrast, the construction of *D. dadantii lpp* length mutants demonstrated that the elongated Lpp+21 variant enhanced the efficacy of OutB attachment to PG, whereas the truncated LppΔ21 variant diminished it. In this context, the length of OutB (OutBΔ22, OutB, or OutB+21) did not affect these trends. It is tempting to suggest that such alterations in the extent of OutB linking to PG are caused by displacements of the PG layer in the periplasm of *D. dadantii lppΔ21* and *lpp+21* mutants. Indeed, previous studies have shown that in *lpp+21* mutants of *E. coli* and *S. enterica*, the PG layer was closer to the IM than in the wild-type strain (36, 37) and exhibited a broader and more diffuse morphology (37). Such a diffuse PG layer may explain the enhanced diffusion of free forms of OutB through the PG mesh of the *D. dadantii lpp+21* mutant (Fig. 3C).

Another reported consequence of the increased periplasm width in the *E. coli lpp+21* mutant was the inactivation of the Rcs signaling pathway (35). In this cellular context, the size of the RcsF sensor lipoprotein has become insufficient to establish contact with the IM sensor IgaA (35). The Rcs signaling pathway is known to activate the expression of genes involved in capsule synthesis (*cps*) and cell division (*ftsZ*), while repressing the transcription of flagellar genes (46, 47). Accordingly, the phenotypes of the *D. dadantii lppΔ21* mutant (mucoid, small, and non-motile cells) (Fig. 3B, S5D and S5E) are indicative of a constitutively activated Rcs system. It seems plausible that, due to the diminished periplasmic width in the *D. dadantii lppΔ21*, the OM-located RcsF remains in continuous contact with the IM sensor IgaA, thereby activating the Rcs cascade. It is likely that the elevated exopolysaccharide production is responsible for the resistance of *D. dadantii lppΔ21* to SDS, despite the apparent alterations in cell morphology (Fig. 3B and S7). It is noteworthy that a search for vancomycin-resistant mutants of *E. coli* has resulted in the isolation of a spontaneous in-frame *lppΔ21* strain with a fourfold increased resistance to the antibiotic (48). It seems plausible that an increased production of exopolysaccharides by the *lppΔ21* mutant could provide generalized protection against desiccation and external agents (49). In contrast, *lpp+21* mutants showed an increased susceptibility to SDS (Fig. 3B and S5D) and to vancomycin (38).

Interestingly, OutB shares a similar topology with TolR, a protein of the trans-envelope Tol-Pal complex. TolR is anchored in the IM by an N-terminal helix that is connected to a periplasmic PG-binding domain by an extensible linker (50, 51). However, the linker of TolR is considerably shorter than that of OutB, 20 *versus* 75 residues. In the resting state, the PG-binding domain of TolR is located near the IM. However, PMF-driven conformational changes in the Tol-Pal complex trigger an extension of the TolR linker so that the PG-binding domain attains and binds the PG layer (50). From this point of view, the length of the OutB linker seems excessive for a rather similar function, i.e. the connection of the IM anchor to the PG-binding domain. The seemingly excessive length of the OutB linker may be related to its role as a scaffolding protein for the secretin OutD. Indeed, the OutB HR domain tightly interacts with the entrance of the large secretin channel that extends into the periplasm (31). A long, flexible linker of OutB may facilitate its contact with the secretin OutD. It is also plausible that the distance between the entrance of the secretin pore and the IM varies during the secretion process, thereby facilitating the recruitment of effector proteins into the secretin channel (52). Consistent with this presumed biological function of OutB, we showed that OutB is expressed in response to envelope stress associated with the plant infection process, conditions essential for the assembly and scaffolding of the OutD secretin pore to the cell envelope.

Site-directed mutagenesis of the Lpp-like box of OutB showed that it can tolerate substantial residue substitutions. For example, the presence of a double positive charge (Lys-Lys) at the C-terminus is not mandatory. The OutB_K220tga variant lacking the C-terminal Lys^220^ was attached to PG by the remaining penultimate Lys^219^, which became the C-terminal residue. Indeed, the truncated Lpp-box of OutB_K220tga (RTTK) is quite similar to those of the Lpp orthologs from *Photobacterium, Vibrio* and *Zobellella*, namely S/RYTK (17). Interestingly, in *E. coli*, over 20% of periplasmic proteins and lipoproteins possess a C-terminal Lys residue (53). Taken together with the apparent absence of a well-conserved Lpp-like consensus, this led us to suggest that some other periplasmic and/or membrane proteins may also be covalently attached to the PG in *D. dadantii* and probably in other proteobacteria. Indeed, the fact that the Lpp-box of OutB is able to drive the covalent linking of a *bona fide* periplasmic protein, BlaM, shows that the nature of the protein itself is not a causal factor. However, as illustrated in this study, the identification of such putative PG-linked proteins by a canonical MS approach may be challenging due to their moderate expression level and the high background noise generated by the highly abundant Lpp.

## Materials and Methods

### Strains, plasmids, growth conditions, and mutagenesis

The bacterial strains and plasmids used in this study are listed in the Table S1. Bacteria were grown in lysogeny broth (LB) at 28°C. If necessary, glucose, glycerol, or arabinose were added at 0.2%. When necessary, antibiotics were supplemented at follows: ampicillin, 50 mg/L; kanamycin, 50 mg/L, and chloramphenicol, 25 mg/L. Site-directed mutagenesis was carried out using the PrimeSTAR Max DNA Polymerase (TaKaRa) with the primers listed in Table S2. The sequences of mutant and amplified genes was checked (Eurofins MWG Operon or Microsynth). *D. dadantii* mutant strains carrying chromosomal *ldt* or *lpp* mutant alleles were constructed by homologous recombination followed by *de novo* transduction of mutant alleles with phage phi-EC2 (54).

### PG purification for Western blotting analysis

*D. dadantii* were grown to late-exponential phase, until an optical density at 600 nm (OD_600_) of about 1.0 in LB supplemented with 0.2% glycerol and appropriate antibiotics. Cells (∼5 × 10^10^) were collected by centrifugation (7,000 g, 5 min), resuspended in 2 mL of 50 mM Tris-HCl, 1 mM EDTA (TE) buffer, then supplemented with 2.5 mL of 10% SDS, boiled at 100°C for 1 h, and stored overnight at 30°C. PG was pelleted by centrifugation (100,000 g, 2 h), resuspended in 4 mL of 2% SDS, boiled for 30 min, and pelleted again (100,000 g, 1 h). Next, PG was resuspended in TE, boiled for 30 min, and collected by centrifugation (20,000 g, 1 h). This washing step was repeated thrice, and PG was resuspended in 0.5 mL of TE. 80 µL of the PG sample was digested for 3 h at 37°C with 4 µg of lysozyme (Sigma-Aldrich) and then, half of the sample was completed with 2 mM MgSO_4_, 0.1 mM ZnSO_4_ and 1 µg of PGRP amidase from *Sitophilus zeamais*, and digested for an additional 1 h at 37°C. The PG samples were loaded on SDS-PAGE immediately after digestion.

### SDS-PAGE and immunoblotting

PG samples digested with lysozyme and PGRP amidase were supplemented with Laemmly loading buffer, boiled for 7 min and undigested PG material was pelleted at 10,000 g for 10 s. The samples were separated in 10% Tris-glycine (for OutB) or 15% tris-tricine (for Lpp) SDS-polyacrylamide gel and transferred to Roti PVDF 0.45 µm membrane (Roth). Blots were probed with rabbit polyclonal antibodies raised against OutB (31) or *D. dadantii* Lpp (this work) at 1:1,500 dilution followed by secondary anti-rabbit goat IgG conjugated to peroxidase (Sigma-Aldrich) at 1:20,000 dilution. Glycoproteins containing *N*-acetylglucosamine (GlcNAc) residues were detected with wheat germ agglutinin, WGA conjugated to peroxidase (Sigma-Aldrich). Chemiluminescence signals generated with Immobilon substrate (Millipore) were detected using Fusion FX imaging system (Vilber) or Hyperfilm (Cytiva) and quantified with EvolutionCapt Edge (Vilber) or with Fiji software.

### Electron Microscopy (EM)

Bacteria were grown in 10 ml of LB at 28°C for 16 h with shaking (120 rpm). Cells were collected by centrifugation (6,000 g, 5 min) and washed in phosphate buffer solution (PBS) pH 7.2 (Sigma-Aldrich). For negative stain EM, the cells were resuspended in 5 ml of PBS and then, adsorbed onto carbon grids (EMSdiasum), and stained with 1.5% uranyl acetate (BHD Chemicals Ltd.) for contrast enhancement. The samples were analyzed with Tecnai Spirit BioTWIN transmission electron microscope at 120 kV. Measurments were made with Radius EM Imaging Software (Emsis). For scanning electron microscopy (SEM), bacteria were grown and washed as described above. Then, cells were fixed overnight in 2.5% glutaraldehyde and postfixed in 1% osmium tetroxide (Agar), and gradually dehydrated in ethanol. Bacteria were mounted onto SEM stubs and coated with gold using a sputter coater spi. The samples were examined and photographed using a Prisma E Scanning Electron Microscope (Thermo Fisher Scientific).

### Mass spectrometry analyses of peptidoglycan strucuture

Peptidoglycan was extracted by the hot SDS procedure and proteins were removed with pronase and trypsin (55). Peptidoglycan was then digested with lysozyme and mutanolysin to obtain soluble disaccharide-peptide fragments. The resulting muropeptides were reduced with NaBH_4_ and separated by *rp*HPLC on a C_18_ column (Hypersil GOLD aQ 250 × 4.6, 3 µm; ThermoFisher) at a flow rate of 1 mL/min. A linear gradient (0% to 100%) was applied between 11.1 min and 105.2 min at 20 °C (buffer A: 0.1% TFA; buffer B: 0.1% TFA, 20% acetonitrile; v/v). Muropeptides were detected by the absorbance at 205 nm. One-mL fractions were individually collected lyophilized, solubilized in 20 µL of water, and stored at -20 °C. Mass spectra were obtained by injecting an aliquot of the fractions (5 μL) into the mass spectrometer (Maxis II ETD, Bruker, France) at a flow rate of 0.1 mL/min (50% acetonitrile, 50% water; v/v acidified with 0.1% formic acid; v/v). Spectra were acquired in the positive mode with a capillary voltage of 3,500V, an *m*/*z* scan range was from 300 to 1,850 at a speed of 2 Hz, as previously described (56). MS/MS spectra were obtained using a collision energy of 50 eV as previously described (56).

### Supporting information

This article contains supporting information (Figures S1-S13, Tables S1 and S2 and Extended Materials and Methods).

## Acknowledgements

We thank Pedro da Silva (BF2I Laboratory, INSA Lyon) for generously providing the PGRP amidase from the weevil *Sitophilus zeamais*. We also thank Guy Condemine and Yvan Rahbe for critical reading of manuscript. This study was supported by funding from the Centre National de la Recherche Scientifique (CNRS) and ANR program SYNERGY_T2SS ANR-19-CE11-0020-01.

