## Supplemental figures S1-S13 and tables S1 and S2 for "Inner membrane protein OutB is covalently attached to peptidoglycan in the γ-proteobacterium *Dickeya dadantii*"

**This file includes**

Figures S1 to S13,

Tables S1 and S2,

Extended Materials and Methods,

SI References

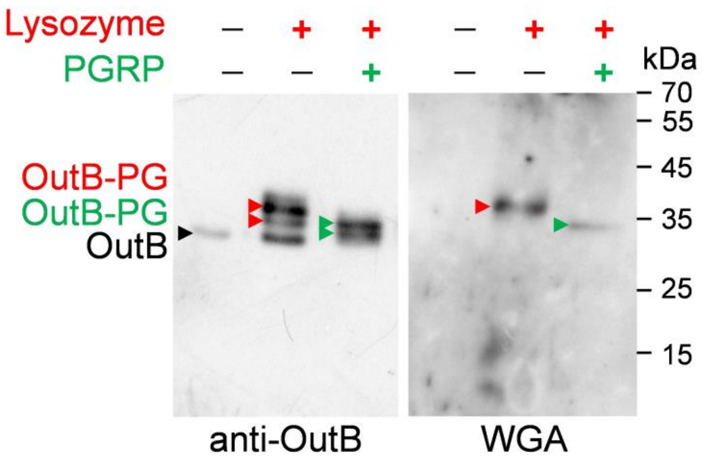


**Figure S1.** Western blot analysis of OutB-PG species linked to muropeptides in *D. dadantii Δlpp* mutant. PG from *D. dadantii Δlpp* mutant expressing *outB* ectopically was digested or not with lysozyme or PGRP amidase and analyzed by Western blot with anti-OutB or WGA. The positions of OutB-PG species generated by lysozyme and PGRP amidase are indicated with red and green arrows, respectively. The position of "free" form of OutB is shown with black arrow.

| Muropeptide | Cross-link | Monoisotopic mass | |
|---|---|---|---|
| | | Calculated | Observed |
| GM$^{Red}$-Tri<br>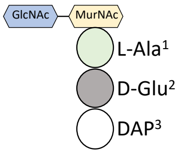 | NA | 870.371 | 870.373 |
| GM$^{Red}$-Tri-Gly<br>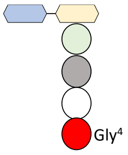 | NA | 927.392 | 927.393 |
| GM$^{Red}$-Tetra<br>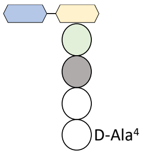 | NA | 941.408 | 941.404 |
| GM$^{Red}$-Tri→Lys<br>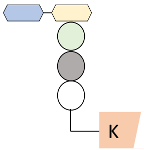 | NA | 998.466 | 998.461 |
| GM$^{Red}$-Tri→Lys-Lys<br>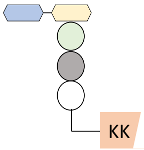 | NA | 1126.561 | 1126.558 |
| GM$^{Red}$-Tetra-GM<br>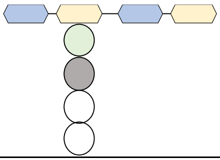 | NA | 1419.588 | 1419.588 |
| GM$^{Red}$-Tri→GM$^{Red}$-Tri<br>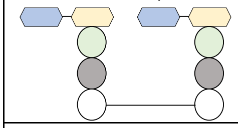 | (3-3) | 1722.731 | 1722.730 |
| GM$^{Red}$-Tetra→GM$^{Red}$-Tri-Gly<br>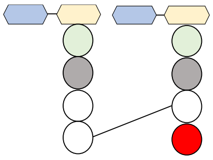 | (4-3) | 1850.789 | 1850.794 |
| GM$^{Red}$-Tetra→GM$^{Red}$-Tri<br>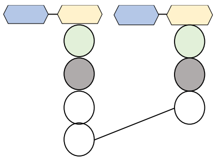 | (4-3) | 1793.768 | 1793.766 |

| | | | |
|---|---|---|---|
| GM$^{Red}$-Tri→GM$^{Red}$-Tetra 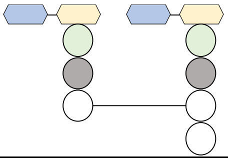 | (3-3) | 1793.768 | 1793.773 |
| GM$^{Red}$-Tetra→GM$^{Red}$-Tri→Lys 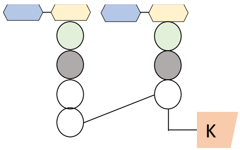 | (4-3) | 1921.863 | 1921.865 |
| GM$^{Red}$-Tetra→GM$^{Red}$-Tetra 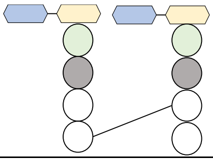 | (4-3) | 1864.805 | 1864.803 |
| GM$^{Red}$-Tri→GM$^{Red}$-Tri→Lys-Lys 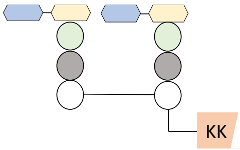 | (3-3) | 1978.921 | 1978.927 |
| GM$^{Red}$-Tetra→GM$^{Red}$-Tri→Lys-Lys 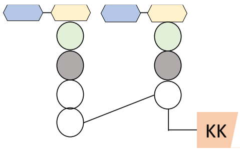 | (4-3) | 2049.958 | 2049.951 |
| GM$^{Red}$-Tetra→GM$^{Anh}$-Tri 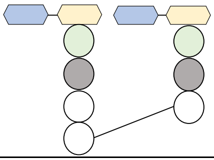 | (4-3) | 1864.805 | 1864.803 |
| GM$^{Red}$-Tetra→GM$^{Anh}$-Tri→Lys-Lys 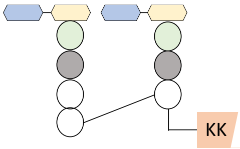 | (4-3) | 2029.931 | 2029.935 |
| GM$^{Red}$-Tetra→GM$^{Anh}$-Tetra 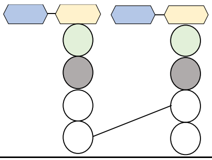 | (4-3) | 1844.779 | 1844.782 |
| GM$^{Red}$-Tetra→GM$^{Red}$-Tetra→GM$^{Red}$-Tri→Lys-Lys 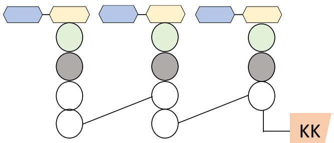 | (4-3) | 2973.355 | 2973.370 |

**Figure S2.** Muropeptide composition of the peptidoglycan of *D. dadantii* strain 3937. Pink trapeze with KK corresponds to the Lys-Lys C-terminal extremity of Lpp and OutB. Abbreviations: Anh, anhydro; Red, reduced; NA, not analyzed.

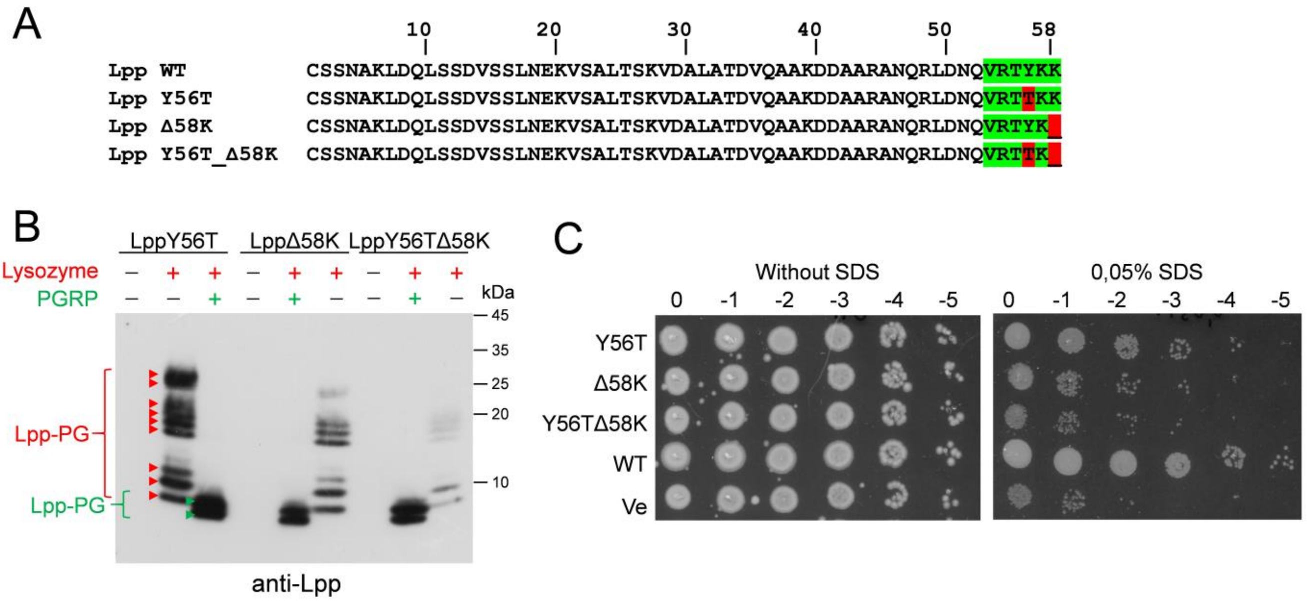


**Figure S3.** Mutagenesis of the Lpp box of $Lpp_{Dd}$. (**A**) Sequence alignment of *D. dadantii* Lpp mutants. The Lpp-box is highlighted in green and the substituted residues are shown in red. (**B**) PG from *D. dadantii Δlpp* mutant strain producing the indicated Lpp variants was digested or not with lysozyme and PGRP amidase and probed by Western blot with anti-Lpp antibodies. Positions of the generated Lpp-PG species are indicated with red and green arrows, respectively. **Please, note that 50-fold more PG material was loaded for this Western blot than on that with the wild-type Lpp shown in Fig. 1C.** (**C**) SDS susceptibility assay with the Lpp variants. Overnight cultures of *D. dadantii Δlpp* ectopically expressing indicated Lpp variants were serially diluted and plated onto LB agar, whether or not containing 0.05% SDS and incubated for 24 h at 28°C.

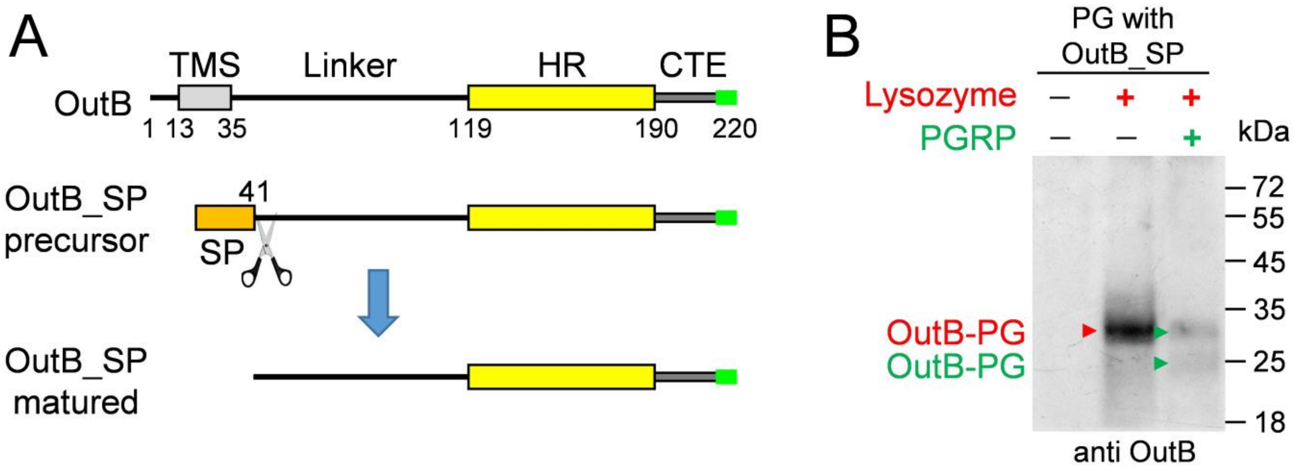


**Figure S4.** The inner membrane anchoring is not essential for attachment of OutB to PG. (**A**) Schematic of OutB_PS variant carrying a cleavable signal peptide (SP) originated from pET-20b in place of the native TMS. (**B**) Western blot analysis of PG purified from *D. dadantii* expressing OutB_PS. PG was digested or not with lysozyme and PGRP amidase and probed with anti-OutB antibodies. Generated OutB-PG adducts are indicated with red and green arrows, respectively.

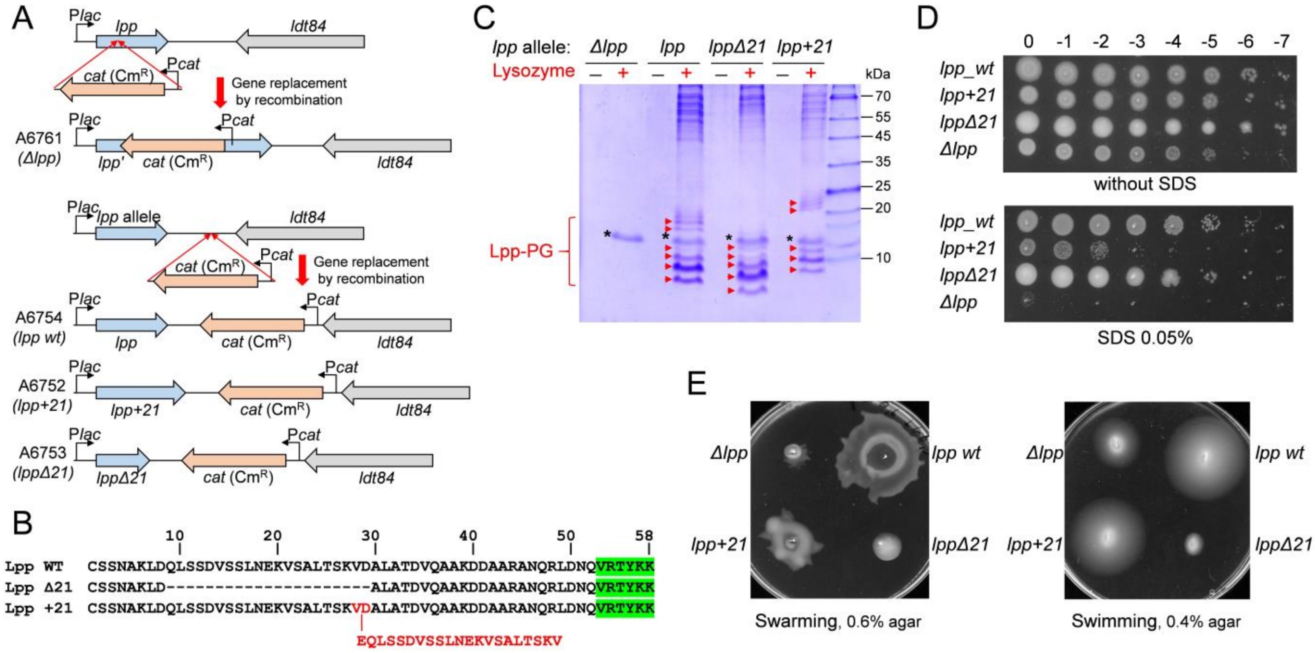


**Figure S5.** Construction and phenotypic assessment of *D. dadantii lpp* mutant strains. (**A**) Construction of *D. dadantii lpp* mutant strains. The mutated *lpp* alleles were introduced into the *D. dadantii* chromosome in place of the wild-type *lpp* gene by homologous recombination. The *cat* ($Cm^R$) gene was used as a selective marker for *de novo* transductions of the mutated *lpp* alleles into the *D. dadantii* wild type strain. (**B**) Sequence alignment of the Lpp length variants. The additional 21-residue sequence of Lpp+21 and its insertion site are shown in red letters. (**C**) PG from *D. dadantii* strains carrying the indicated (on top) *lpp* alleles was purified, digested or not with lysozyme and analyzed by SDS-PAGE. Lpp-PG species are shown with red arrows and lysozyme is noted with an asterisk. (**D**) SDS susceptibility assay. Overnight cultures were serially diluted and plated onto LB agar, whether or not containing 0.05% SDS and incubated for 24 h at 28°C. (**E**) Swarming (left) and swimming (right) motility assays. ~$10^6$ of overnight grown cells were deposited into the soft LB agar (0.6 or 0.4%, respectively) and cultivated at 28°C for 14 h.

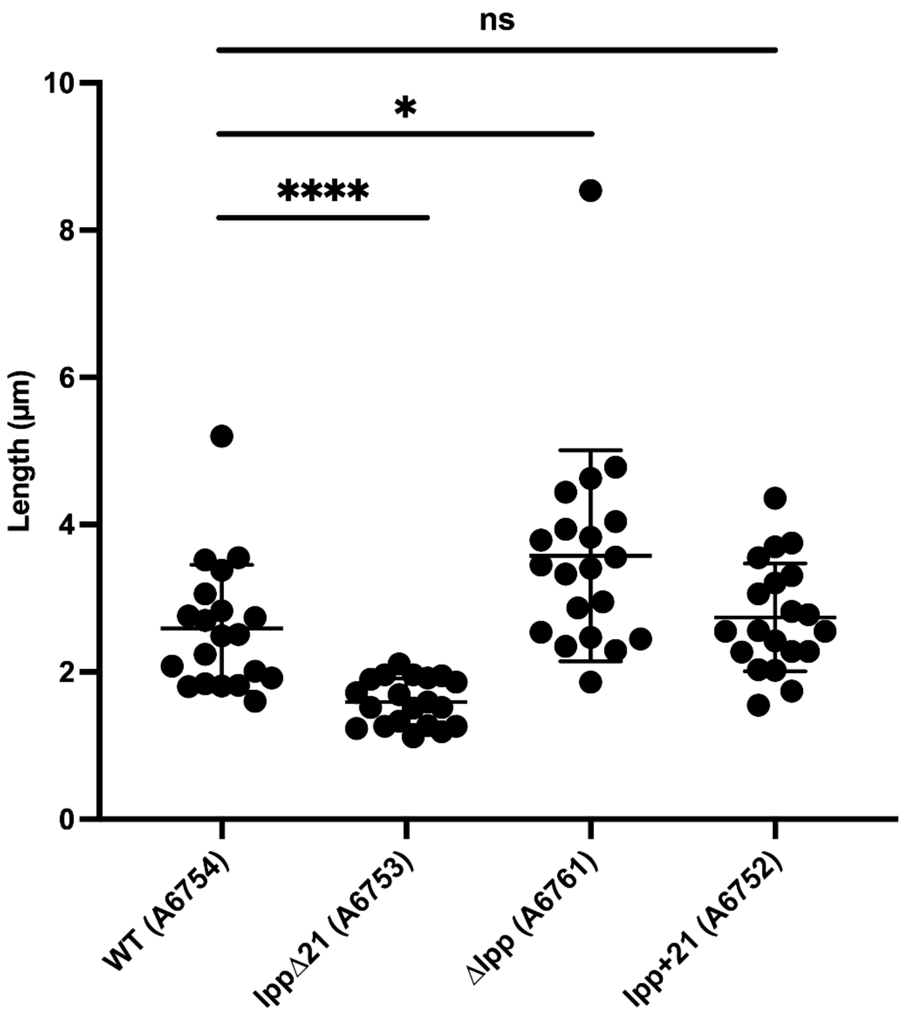


**Figure S6.** Length of *D. dadantii lpp* mutant cells observed in negative stain EM micrographs. For each strain, 20 micrographs with 20 cells were randomly selected. All the collected values, including outliers, were taken for analysis. The data were analyzed with PRISM software using two-sample *t*-test by comparing each mutant to the wild-type strain. Dots represent the length value of each cell. Mean and SEM are shown with horizontal bars. **** and * denote statistically significant differences between mean values compared to the WT strain A6754, with *P* values <0,0001 and 0,0319, respectively.

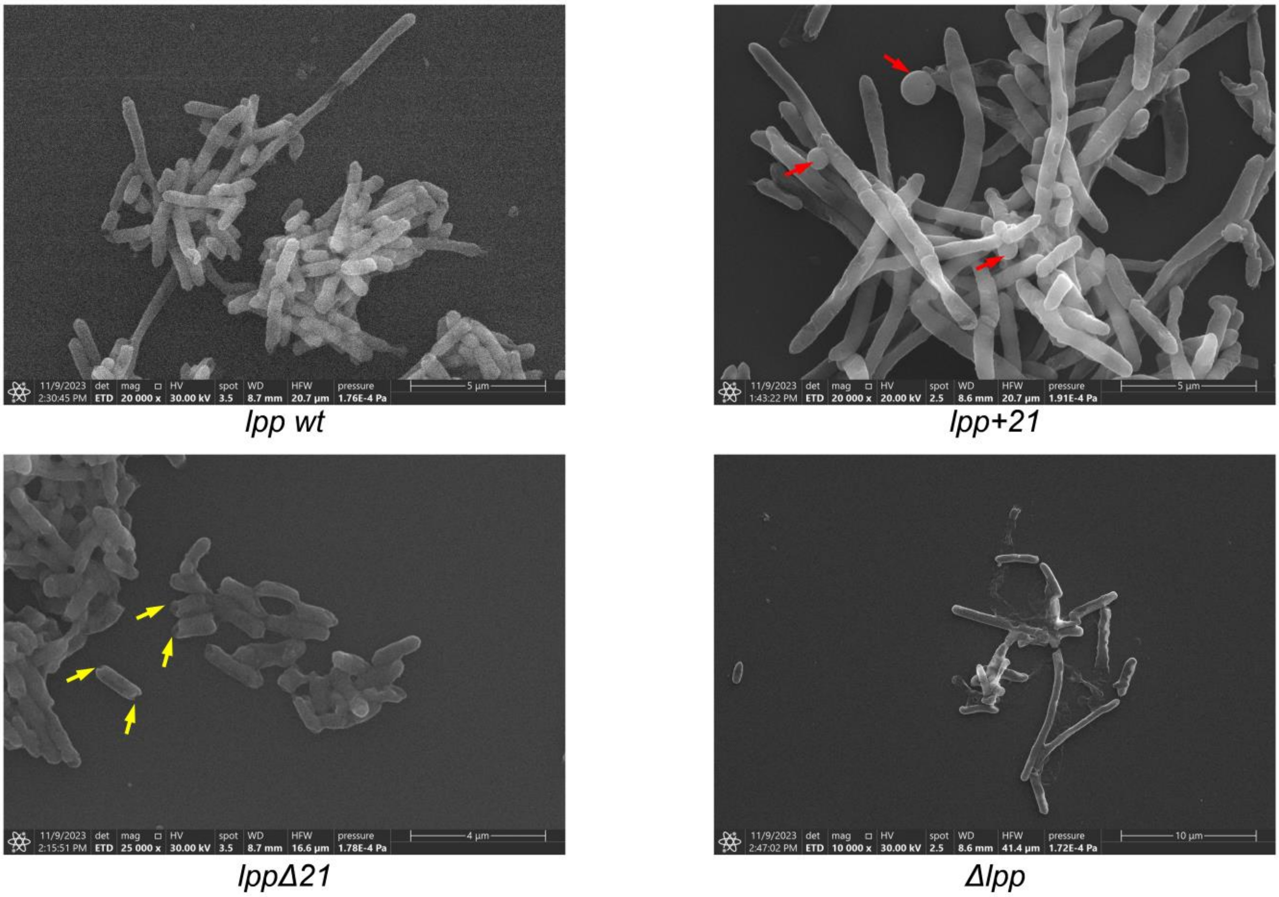


**Figure S7.** Scanning Electron Microscopy of the indicated *D. dadantii lpp* mutant strains. The blebs in the *lpp+21* strain are shown with red arrows and the cavities at the cell poles of *Δlpp* mutant are indicated with yellow arrows.

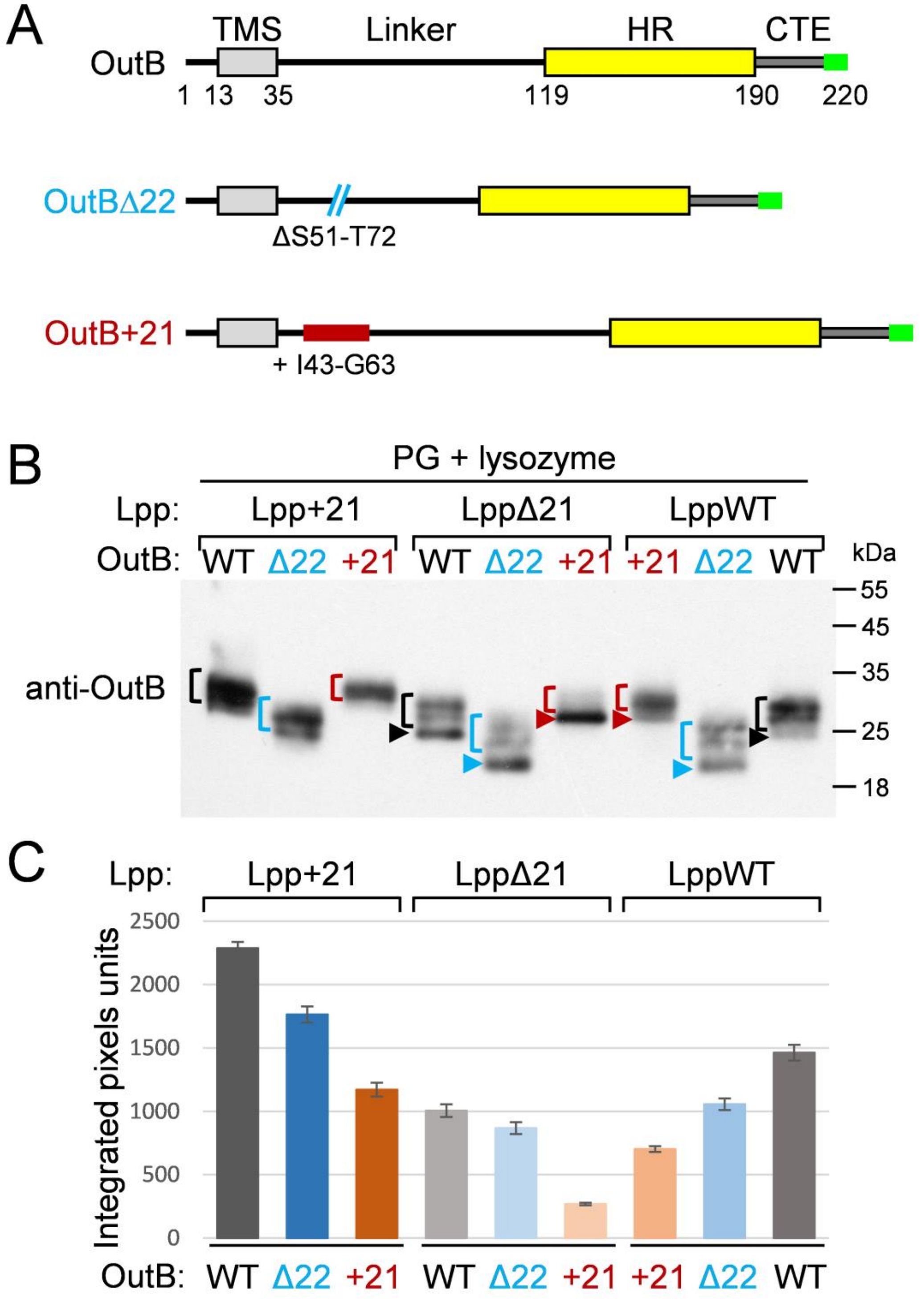


**Figure S8.** Lengthening of Lpp improves attachment of OutB length variants to PG. (**A**) Schematic of OutB length variants. The regions deleted (Ser$^{51}$ to Thr$^{72}$) or inserted (Ile$^{43}$ to Gly$^{63}$) in OutBΔ22 and OutB+21, respectively, are indicated. (**B**) Representative immunoblotting analysis of PG purified from *D. dadantii* WT, *lppΔ21*, and *lpp+21* strains producing either OutBwt, OutBΔ22, or OutB+21 (nine combinations indicated on the top of panel). Only lysozyme-digested samples are shown. "Free" forms of OutBwt, OutBΔ22 and OutB+21 are indicated with black, blue and dark red arrowheads, respectively. OutB-PG species linked to muropeptides are shown with brackets of the same colors. (**C**) Muropeptide-linked specie of each OutB variant (shown with brackets in panel B) were quantified with EvolutionCapt Edge Software (Viber Lourmat) according to the pixel intensity and area of each protein band. The data are from the replicate shown in panel B. The bar heights show the relative amount of muropeptide-linked species for each OutB-length variant in each *D. dadantii lpp*-length mutant line (as they are shown with brackets in panel B). The bars are positioned below the corresponding gel lines in panel B.

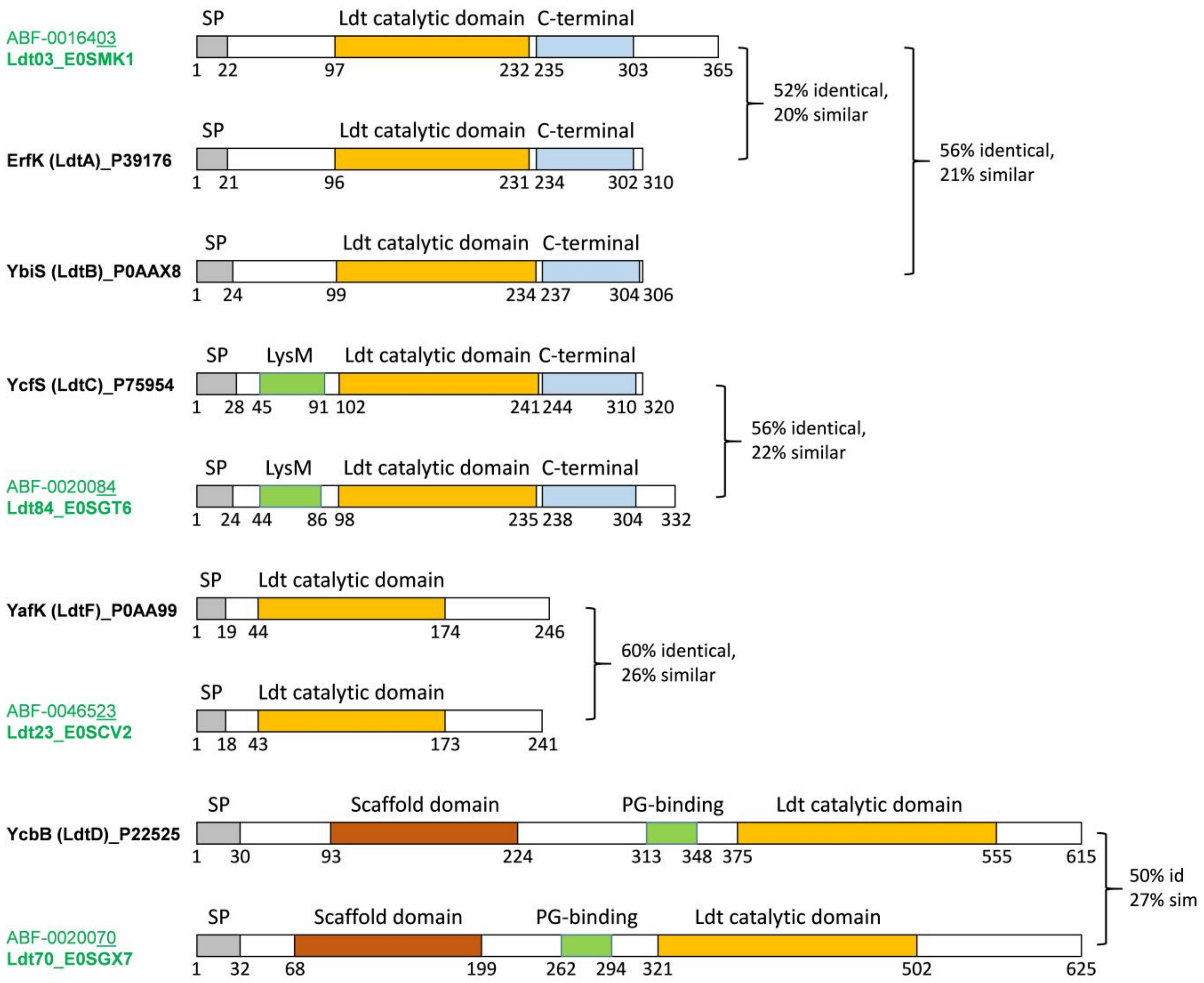


**Figure S9.** Domain organization of L,D-transpeptidases of *D. dadantii* compared with those of *E. coli*. Domain occurrence and their positions are from InterPro (10.1093/nar/gkac993). The names of Ldts are followed with their UniProtKB codes. The names of *D. dadantii* Ldts are in green and those of *E. coli* are in black. The identity and similarity levels of orthologous Ldts are indicated.

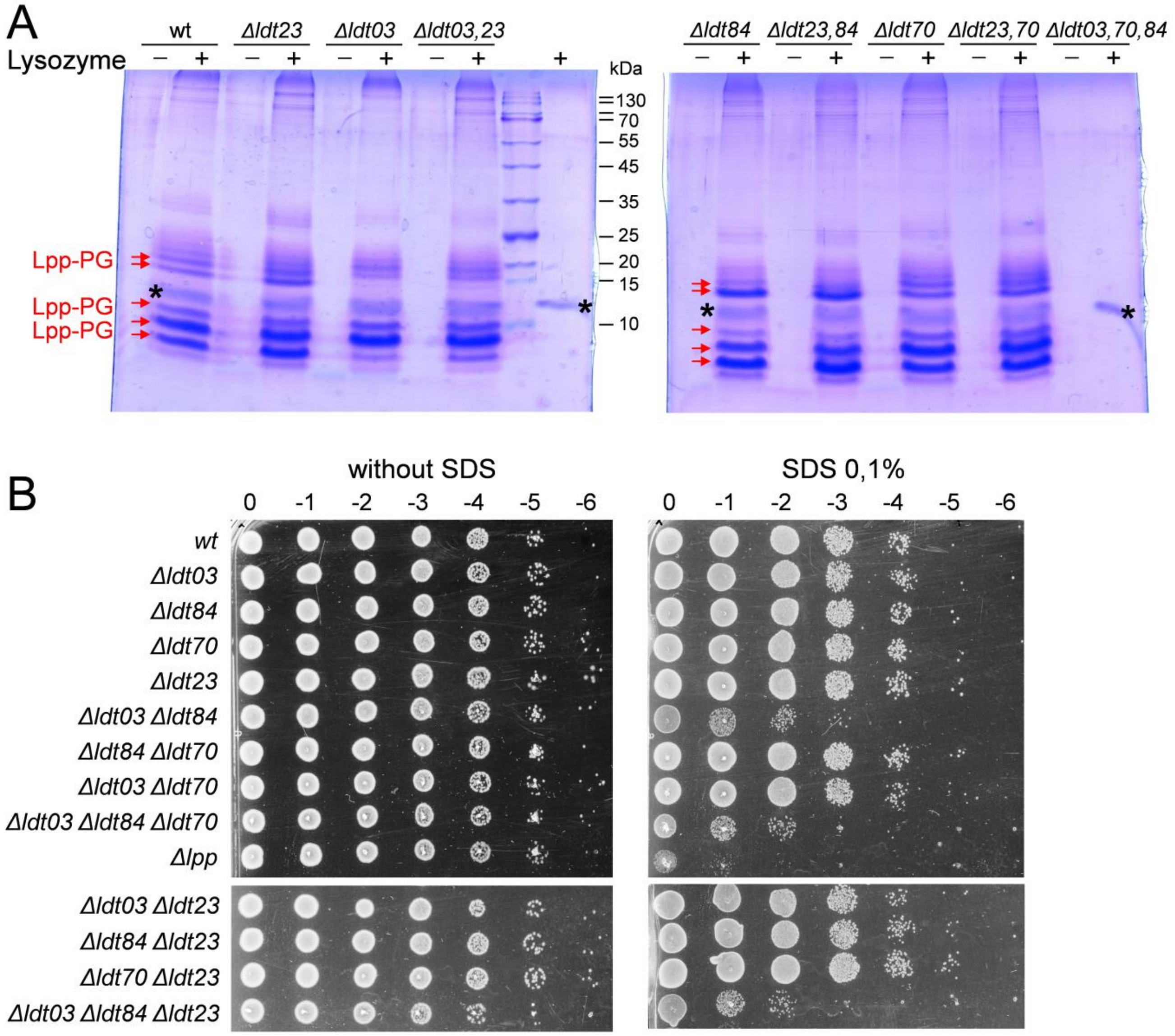


**Figure S10.** Characterization of *D. dadantii Δldt* mutant strains. (**A**) SDS-PAGE analysis of the protein content of PG purified from *D. dadantii Δldt* mutant strains. PG was digested or not with lysozyme. Lpp-PG adducts generated by lysozyme are indicated with red arrows and lysozyme position is shown with an asterisk. (**B**) SDS susceptibility assay with *D. dadantii Δldt* mutant strains. Overnight cultures were serially diluted and plated onto LB agar, whether or not containing 0.1 % SDS and incubated for 24 h at 28°C.

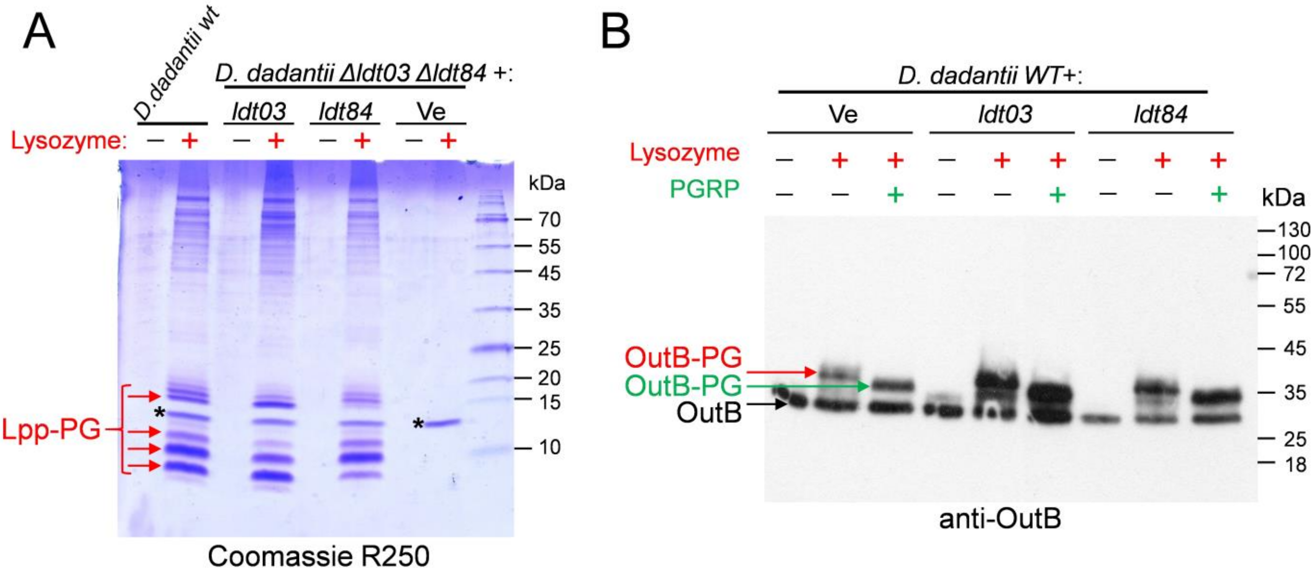


**Figure S11.** Ldt03 and Ldt84 generate different muropeptide linked patterns of Lpp and OutB. (**A**) Comparative analysis of Lpp-muropeptide patterns generated by Ldt03 and Ldt84. PG from *D. dadantii* wild type or *Δldt03 Δldt84* double mutant ectopically expressing either *ldt03,* or *ldt84,* or empty vector (Ve) was digested or not with lysozyme and analyzed by SDS-PAGE. Lpp-PG adducts are indicated with red arrows. (**B**) Ectopic expression of *ldt03* or *ldt84* increased covalent attachment of OutB to PG in the *D. dadantii* WT strain. PG from *D. dadantii* WT carrying either empty plasmid (Ve) or that with *ldt03* or *ldt84* was digested or not with lysozyme and PGRP and analyzed by Western blot with anti-OutB antibodies. The positions of "free" OutB and OutB-PG adducts generated by lysozyme and PGRP are indicated with black, red and green arrows, respectively.

A

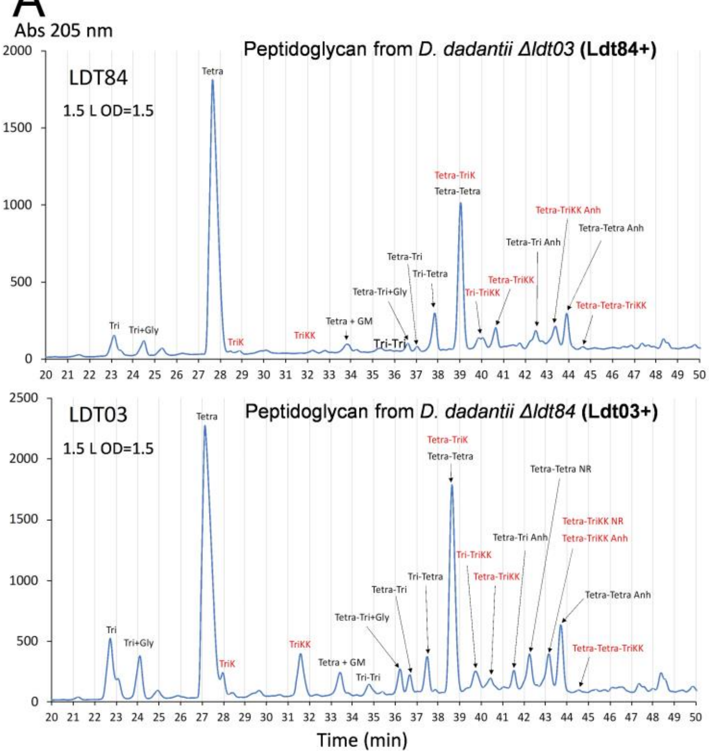


B

**Relative abundance of muropeptides resulting from the PG-anchoring of Lpp**

| | Relative intensity (%) in strains lacking *ldt03* or *ldt84* | | | |
|---|---|---|---|---|
| | *Δldt03* (Ldt84+) | | *Δldt84* (Ldt03+) | |
| **Replicate** | **Tri→KK** | **Tetra→Tri→KK** | **Tri→KK** | **Tetra→Tri→KK** |
| 1 | 16.7 | 83.3 | 86.0 | 14.0 |
| 2 | 33.4 | 66.6 | 73.6 | 26.4 |
| 3 | 33.8 | 66.2 | 77.8 | 22.2 |
| Mean | 28.0 | 72.0 | 79.1 | 20.9 |
| Standard deviation | 9.8 | 9.8 | 6.3 | 6.3 |

C

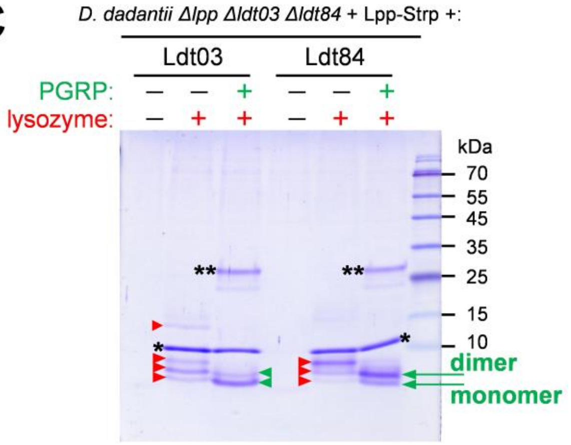


**Figure S12.** Ldt03 and Ldt84 preferentially attach Lpp to monomeric and dimeric peptide stems, respectively. (**A**) *rp*HPLC profile analysis of muropeptides generated from *D. dadantii Δldt03* (Ldt84+, upper panel) and *D. dadantii Δldt84* (Ldt03+, lower panel). Bacteria were grown until late exponential phase ($DO_{600}$ ~2.0) and PG was extracted by the hot SDS-procedure, digested with pronase, trypsin, and muramidases. The resulting muropeptides were reduced with $NaBH_4$, separated by *rp*HPLC and identified by MS. The names of muropeptides plotted on the pics correspond to those in Fig. S2. Muropeptides bearing Lys or Lys-Lys are in red. (**B**) Relative abundance of the Tri→KK monomer and Tetra→Tri→KK 4→3 cross-linked dimer containing the Lys-Lys motif in the PG extracted either from *D. dadantii Δldt03* carrying a chromosomic copy of the *ldt84* gene (Ldt84) or from *D. dadantii Δldt84* carrying a chromosomic copy of *ldt03* (Ldt03). Values are the ion current intensity of corresponding analytes. The respective muropeptide patterns are shown in panel A. (**C**) Comparative assessment of Lpp-muropeptide patterns generated by Ldt03 and Ldt84. PG from *D. dadantii Δlpp Δldt03 Δldt84* triple mutant ectopically expressing *lpp-Strep* together with either *ldt03* or *ldt84* was digested or not with lysozyme and PGRP amidase and analyzed by SDS-PAGE. Lpp-Strep-PG species linked to muropeptides generated by lysozyme and PGRP are indicated with red and green arrowheads, respectively.

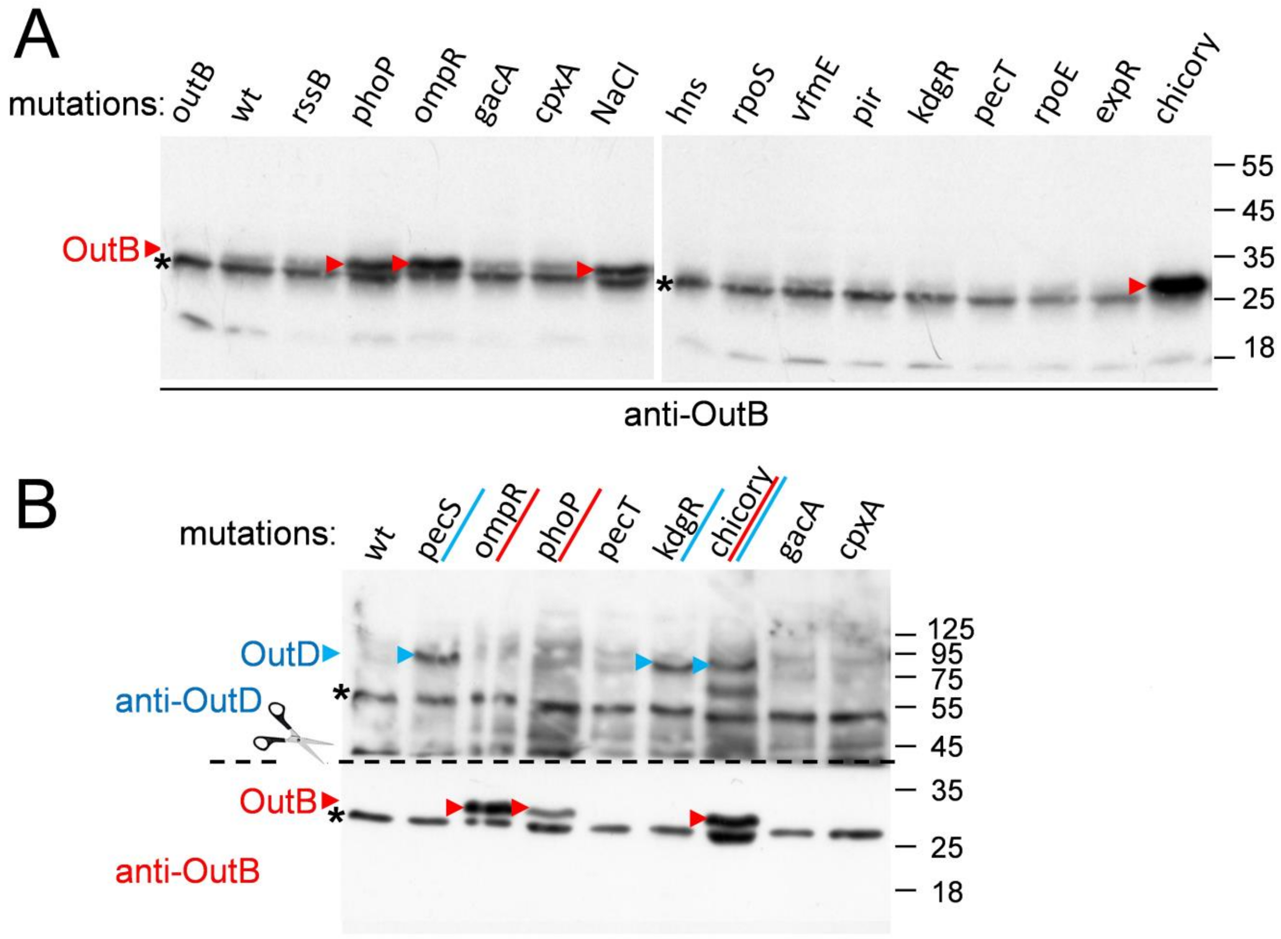


**Figure S13. Production of OutB in *D. dadantii* regulatory mutants.** (**A**) Cell extracts from the indicated *D. dadantii* mutants were separated on SDS-PAGE and analyzed by Western with anti-OutB antibodies. Bacteria were grown in LB supplemented with 0.2 % glycerol at 28°C for 14 h. An equivalent of $10^7$ cells was loaded onto each line. In "chicory", bacteria were grown in chicory leaves. In "NaCl", the medium was supplemented with 0.2 M NaCl. OutB is shown with red arrows. Non-specific species are noted with an asterisk. (**B**) Expression of *outB* and *outD* is not co-regulated by the same regulatory genes. Cell extracts from *D. dadantii* regulatory mutants were prepared and separated on SDS-PAGE as in panel A. After transfer, the upper and lower parts of the blot were probed separately, with anti-OutD or anti-OutB, respectively. The positions of OutD and OutB are shown with blue and red arrows, respectively. Non-specific species are noted with an asterisk.

**Table S1.** Bacterial strains and plasmids used in this study.

| Strains/Plasmids | Genotype/phenotype | Reference |
|---|---|---|
| *Escherichia coli* | | |
| NM522 | *supE thi-1* Δ(*lac-proAB*) Δ(*mcrB-hsdSM*)*5* ($r_K^-$ $m_K^+$) [F′ *proAB lacI*$^q$*Z*Δ*M15*] | NEB |
| *Dickeya dadantii* 3937 | | |
| A5652 | wild type | laboratory collection |
| A5654 | *outB::uidA-nptI* (Km$^R$) | (1) |
| A6259 | *ldt03::tetA* (Tc$^R$) | This work |
| A6260 | *ldt84::aadA* (Sp$^R$/Sm$^R$) | This work |
| A6261 | *ldt70::cat* (Cm$^R$) | This work |
| A6263 | *ldt03::tetA ldt84::aadA* (Tc$^R$ Sp$^R$/Sm$^R$) | This work |
| A6264 | *ldt70::cat ldt84::aadA* (Cm$^R$ Sp$^R$/Sm$^R$) | This work |
| A6266 | *ldt03::tetA ldt84::aadA ldt70::cat* (Tc$^R$ Sp$^R$/Sm$^R$ Cm$^R$) | This work |
| A6267 | *ldt03::tetA ldt70::cat* (Tc$^R$ Cm$^R$) | This work |
| A6300 | *ldt03::tetA ldt84::aadA ldt23::aacC* (Tc$^R$ Sp$^R$/Sm$^R$ Gm$^R$) | This work |
| A6301 | *ldt23::aacC* (Gm$^R$) | This work |
| A6302 | *ldt03::tetA ldt23::aacC* (Tc$^R$ Gm$^R$) | This work |
| A6303 | *ldt84::aadA ldt23::aacC* (Sp$^R$/Sm$^R$ Gm$^R$) | This work |
| A6304 | *ldt70::cat ldt23::aacC* (Gm$^R$ Cm$^R$) | This work |
| A6589 | *ldt03::tetA ldt84::aadA lpp::cat* (Tc$^R$ Sp$^R$/Sm$^R$ Cm$^R$) | This work |
| A6752 | *lpp+21 cat* (Cm$^R$) inserted between *lpp+21* and *ldt84* | This work |
| A6753 | *lpp*Δ*21 cat* (Cm$^R$) inserted between *lpp*Δ*21* and *ldt84* | This work |
| A6754 | *lpp_wt cat* (Cm$^R$) inserted between *lpp* and *ldt84* | This work |
| A6761 | *lpp::cat* (Cm$^R$) | This work |
| **Plasmids** | | |
| pGEM-T | P*lac*, P*T7pol*, *blaM* (Ap$^R$), ColE1 origin | Promega |
| pBAD33 | $P_{BAD}$, *cat* (Cm$^R$), p15A origin | (2) |
| pBS | Bluescript KS+, *blaM* (Ap$^R$), ColE1 origin | Stratagene |
| pBS-Km | Bluescript KS+, *neo* (Km$^R$), ColE1 origin | This work |
| pET-20b(+) | P*T7pol*, coding PelB signal peptide and 6His, *blaM* (Ap$^R$) | Novagen |
| BS | pGEM-T carrying *outB-outS* 1.6 kb fragment | This work |
| pBAD_Ldt03 | pBAD33 carrying *ldt03* under $P_{BAD}$ | This work |
| pBAD_Ldt23 | pBAD33 carrying *ldt23* under $P_{BAD}$ | This work |
| pBAD_Ldt70 | pBAD33 carrying *ldt70* under $P_{BAD}$ | This work |
| pBAD_Ldt84 | pBAD33 carrying *ldt84* under $P_{BAD}$ | This work |
| pBS_Ldt03 | pBS-Km carrying *ldt03* under P*lacZ* | This work |
| pBS_Ldt23 | pBS-Km carrying *ldt23* under P*lacZ* | This work |
| pBS_Ldt70 | pBS-Km carrying *ldt70* under P*lacZ* | This work |
| pBS_Ldt84 | pBS-Km carrying *ldt84* under P*lacZ* | This work |
| pGEM-T_Lpp | pGEM-T carrying the *lpp* gene of *D. dadantii* under P*lacZ* | This work |
| pBAD_Lpp | pBAD33 carrying the *lpp* gene of *D. dadantii* under $P_{BAD}$ | This work |

**Table S2.** Primers used in this study.

| Primer | Nucleotide sequence (5'-3')[b] | Generated mutation[c] or cloned gene |
|---|---|---|
| OuB_T218Y[a] | cggagcaaaccgtcaggaca**tac**aagaaatgacacagcaactgc | OutB_T218Y |
| OuB_K219TGA[a] | gcaaaccgtcaggacaacg**tga**aaatgacacagcaactgcac | OutB_K219tga |
| OuB_K220tga | gaag**tg**atgacacagcaactgcacatc | OutB_K220tga |
| ROuB_K220tga | gtcat**ca**cttcgttgtcctgacggtttg | OutB_K220tga |
| OuB_Msc[a] | gattcgttgcaggattggcc**a**ggcgggaaaccgggcgatg | OutB_*Msc*I over P191-G192 |
| Bla_Msc[a] | cctcactgattaagcattgg**cc**actgtcagaccaagtttactc | BlaM_*Msc*I over W286 |
| BladltB | gacggtttgcggtttcccgcctggccaatg | OutB_ΔG196-E212=OutBΔC |
| RBladltB | gggaaaccgcaaaccgtcaggacaacgaag | OutB_ΔG196-E212=OutBΔC |
| OuBdltlink | ccgctgtcgactgcggatgaaaaccatcc | OutB_ΔS51-T72 = OutBΔ22 |
| ROuBdltlink | catccgcagtcgacagcggaatggggctg | OutB_ΔS51-T72 = OutBΔ22 |
| OuB_Sfo[a] | ccgcctaccaaagtgggc**gcc**gttccggcatcaacagcgg | OutB_*Sfo*I over G63-M64 |
| For_6403 | gtaccggataacaacaattcc | *ldt03* |
| Rev_6403 | gcagcgtaagctgccctgtg | *ldt03* |
| For20084 | gtaaccctgtgttgcgctctg | *ldt84* |
| Rev20084 | gatgtcagaacgcccgctgatg | *ldt84* |
| For_20070 | cgcgatgtgataaaagcagtg | *ldt70* |
| Rev_20070 | gcccgtcgatgcaacccgag | *ldt70* |
| F46523 | cctgcggcttgaaggatgtc | *ldt23* |
| RC_46523 | ctcagccgttgaccggcaac | *ldt23* |
| F_LppRBS | caatttagagggtattaataatg | *lpp* |
| Rev_Lpp | tggcgcacaaagtgcgccat | *lpp* |
| Lpp_Xho[a] | ggttgctccagcaatgctaaact**c**ga**g**cagctgtcttctgacgtttcttc | Lpp_*Xho*I over L7-D8[d] |
| Lpp_SalI[a] | gctctgaccagcaaagt**c**gacgctctggctaccg | Lpp_*Sal*I over V28-D29[d] |
| LppK57R[a] | caaccaggttcgtacttaca**g**gaagtaagaactggttgaatg | Lpp_K57R[d] |
| LppY56T[a] | cctggacaaccaggttcgtact**ac**caagaagtaagaactggttg | Lpp_Y56T[d] |
| Lpp57 | caag**t**agtaagaactggttgaatgaaaaatg | Lpp_Δ58K[d] |
| RLpp57 | cttact**a**cttgtaagtacgaacctggttgtcc | Lpp_Δ58K[d] |
| Lpp_dlt58_Y56T[b] | gtact**ac**caagtagtaagaactggttgaatg | Lpp_Y56T_Δ58K[d] |
| RLpp_dlt58_Y56T[b] | cttg**gt**agtacgaacctggttgtccagg | Lpp_Y56T_Δ58K[d] |

[a] For each primers used in site directed mutagenesis, another primer with reverse complementary sequence was used (not shown).

[b] *lppΔ58K* was used as template.

[c] Mutated or introduced bases are in bold.

[d] The Lpp residue numbering is this for the matured, signal peptide-less Lpp.

**Plasmid and strain construction.**

To generate OutB+21, a *Sfo*I site was introduced into the *outB* sequence covering the codons of G63 and M64. Next, the *Eco*RI-*Sfo*I *outB* fragment encoding the residues M1 to G63 was fused to the *Eco*RV-*Eco*RI *outB* fragment encoding the residues I43 to K220. In this way, in OutB+21, the sequence I43 to G63 was repeated twice. OutBΔ22 variant missing residues S51 to T72 was generated by PCR using OuBdltlink and ROuBdltlink primers. OutBΔC variant missing residues G196-E212 was generated by PCR using BladltB and RBladltB primers. To generate BlaM-CTE fusion in pGEM-T vector, an *Msc*I site was introduced into the *blaM* sequence covering the W286 codon. Next the *Ssp*I-*Msc*I *blaM* fragment from pGEM-T was inserted into *Sma*I-*Msc*I sites of the BS plasmid in the place of *outB* sequence encoding residues M1 to W190. In this way, full-length BlaM was fused to P191-K220 fragment of OutB.

To generate LppΔ21, an *Xho*I and a *Sal*I sites were introduced into the *lpp* sequence, covering respectively, the codons of L7-D8 and V28-D29 of the mature, signal peptide-less Lpp. Next, the *Xho*I-*Sal*I *lpp* fragment encoding residues D8 to V28 was deleted, producing *lppΔ21*. To generate Lpp+21, the *Xho*I-*Sac*II *lpp* fragment encoding residues Q9 to K58 was fused to the *Sal*I-*Sac*II *lpp* fragment encoding residues C1 to E29. In this way, in Lpp+21, the sequence Q9 to E29 was repeated twice.

Mutant *D. dadantii* strains carrying chromosomal mutations in *ldt*, *lpp* or *outB* genes were generated by by homologous recombination. To this end, a *tetA* ($Tc^R$) cartridge from pHP45Ω-Tc plasmid was introduced into the unique *Eco*RI site of *ldt03*, an *aadA* ($Sp^R$/$Sm^R$) cartridge from pHP45Ω-Sm plasmid was introduced into the unique *Bam*HI site of *ldt84*; an *aacC* ($Gm^R$) cartridge from p34S-Gm plasmid was introduced into the unique *Bam*HI site of *ldt23*, and a *cat* ($Cm^R$) cartridge from pCKC15 plasmid was introduced into the unique *Age*I site of *ldt70* and into the unique *Hinc*II site of *lpp* (3-5).

Construction of *Δlpp*, *lppΔ21* and *lpp+21* mutant strains is shown in Fig. S5. The *cat* ($Cm^R$) gene, inserted between the *lpp* and *ldt84* genes was used as a selective marker for *de novo* transductions of the mutated *lpp* alleles into the *D. dadantii* wild type.
